# Multiomic Evidence for a Unified Model of Alzheimer’s Disease Etiology Linking Microglial Flux Capacity and Astrocyte-Neuron Metabolic Breakdown

**DOI:** 10.1101/2024.07.23.604835

**Authors:** Tom Paterson, Jennifer Rohrs, Timothy J. Hohman, Mark Mapstone, Peter J. Meikle, Rima Kaddurah-Daouk, Allan I. Levey, Leroy Hood, Alzheimer’s Disease Neuroimaging Initiative, Alzheimer Disease Metabolomics Consortium, Cory C. Funk

**Author notes:** Data used in preparation of this article were generated by Alzheimer Disease Metabolomics Consortium (ADMC) using samples obtained from the Alzheimer’s Disease Neuroimaging Initiative. (ADNI) database (adni.loni.usc.edu). As such, the investigators within the ADNI contributed to the design and implementation of ADNI and/or provided data but did not participate in analysis or writing of this report. A complete listing of ADNI investigators can be found at: http://adni.loni.usc.edu/wp-content/uploads/how_to_apply/ADNI_Acknowledgement_List.pdf.

## Abstract

Age and APOE genotype are the strongest known risk factors for late-onset Alzheimer’s disease (AD), but the mechanisms linking them to neuronal loss remain incompletely defined. Using multiomic data from the Alzheimer’s Disease Neuroimaging Initiative (ADNI), we propose a unified hypothesis in which two interdependent failure modes—saturation of microglial lipid flux capacity and disruption of the astrocyte–neuron lactate shuttle (ANLS) due to excess astrocytic membrane cholesterol—drive disease progression upstream of amyloid and tau pathology. Stratifying participants by cognitive score quartiles, we find consistent associations linking impaired lipid clearance, metabolic stress, and genetic variants regulating cholesterol handling. These processes appear to reinforce each other, resulting in accelerating neurodegeneration. Our hypothesis reframes AD as a systems-level collapse in metabolic coordination, rather than a purely linear pathological cascade. These insights emerged during the development of digital twin models for personalized interventions, highlighting the power of systems approaches to reveal hidden drivers of neurodegeneration.

## Introduction

Alzheimer’s disease (AD) was first described by Alois Alzheimer in 1907, who identified hallmark features including amyloid plaques, neurofibrillary tangles, and lipid inclusions— structures he termed “adipose saccules” and which are now recognized as lipid droplets (LDs)^1^. Today, age and APOE genotype, particularly the APOE4 allele, remain the most robust risk factors for late-onset AD^2^. Genome-wide association studies (GWAS) have expanded this picture, implicating genes involved in lipid metabolism, glial clearance, and endolysosomal trafficking^3,4^. Yet a unified mechanistic model linking these processes to progressive neurodegeneration remains lacking.

In the healthy brain, three major mechanisms regulate cholesterol efflux. The first is reverse cholesterol transport, in which astrocyte-derived APOE-containing lipoproteins mediate cholesterol redistribution and clearance via ABCA1 and ABCG1 transporters^5,6^. The second is neuronal conversion of cholesterol into 24S-hydroxycholesterol (24-OHC), which crosses the blood-brain barrier (BBB) and is metabolized in the liver^7,8^. The third pathway, particularly active in glial cells, is the conversion of cholesterol into 27-hydroxycholesterol (27-OHC), a locally acting oxysterol that activates liver X receptor alpha (LXRA) in the CNS^9,10^. Activation of LXRA represses SREBP2, the master regulator of de novo cholesterol synthesis, forming a feedback loop to maintain cellular cholesterol balance^11,12^.

Lipid droplets act as intracellular buffers of cholesterol and are known to vary with physiological state, including diurnal rhythms^13–16^. However, in AD, this system becomes overwhelmed. With age and increased myelin turnover, glia are exposed to rising levels of neuronal membrane debris, challenging their capacity to traffic cholesterol efficiently. The sheer volume of excess cholesterol saturates lipoprotein particles, causing lipid droplets to enlarge and persist^17^. As lipid droplets grow, they lose access to mitochondria, impairing the synthesis of 27-OHC. This disrupts LXRA activation, lifting repression on SREBP2, and allowing unchecked de novo cholesterol synthesis even in an already overloaded environment^11,12^. This breakdown in astrocytic cholesterol homeostasis impairs their ability to metabolically support neurons and contributes to early Alzheimer’s pathogenesis^18^.

To investigate how these disruptions contribute to cognitive decline, we analyzed baseline multiomic data from the Alzheimer’s Disease Neuroimaging Initiative (ADNI), stratifying participants into quartiles based on ADAS13 cognitive scores^18^. Our approach enriched for common pathological signatures independent of strict diagnostic categories. Across these quartiles, we identified two interdependent processes that appear to drive disease progression: saturation of microglial lipid flux capacity, and cholesterol-induced impairment of the astrocyte– neuron lactate shuttle (ANLS). These feedback loops emerge upstream of amyloid and tau pathology and show evidence of mutual reinforcement.

Our unified model integrates lipidomic, proteomic, and genetic data, linking impaired lipid handling with neuronal energetic failure. The hypothesis arose during the development of a digital twin model for personalized interventions, where failure to accurately predict biomarker trajectories pointed toward missing biological constraints—specifically, glial processing-capacity limits. Together, these findings offer a new framework for understanding late-onset AD as a systems-level breakdown of metabolic homeostasis.

## RESULTS

### Data Selection/Organization

To investigate how glial lipid processing and neuronal metabolic support vary with cognitive status, we analyzed baseline multiomic data from ADNI. Participants were stratified into quartiles based on ADAS13 cognitive scores^19^, providing finer resolution across the cognitive spectrum independent of clinical diagnosis (**Figure 1C**).

**Figure 1.**
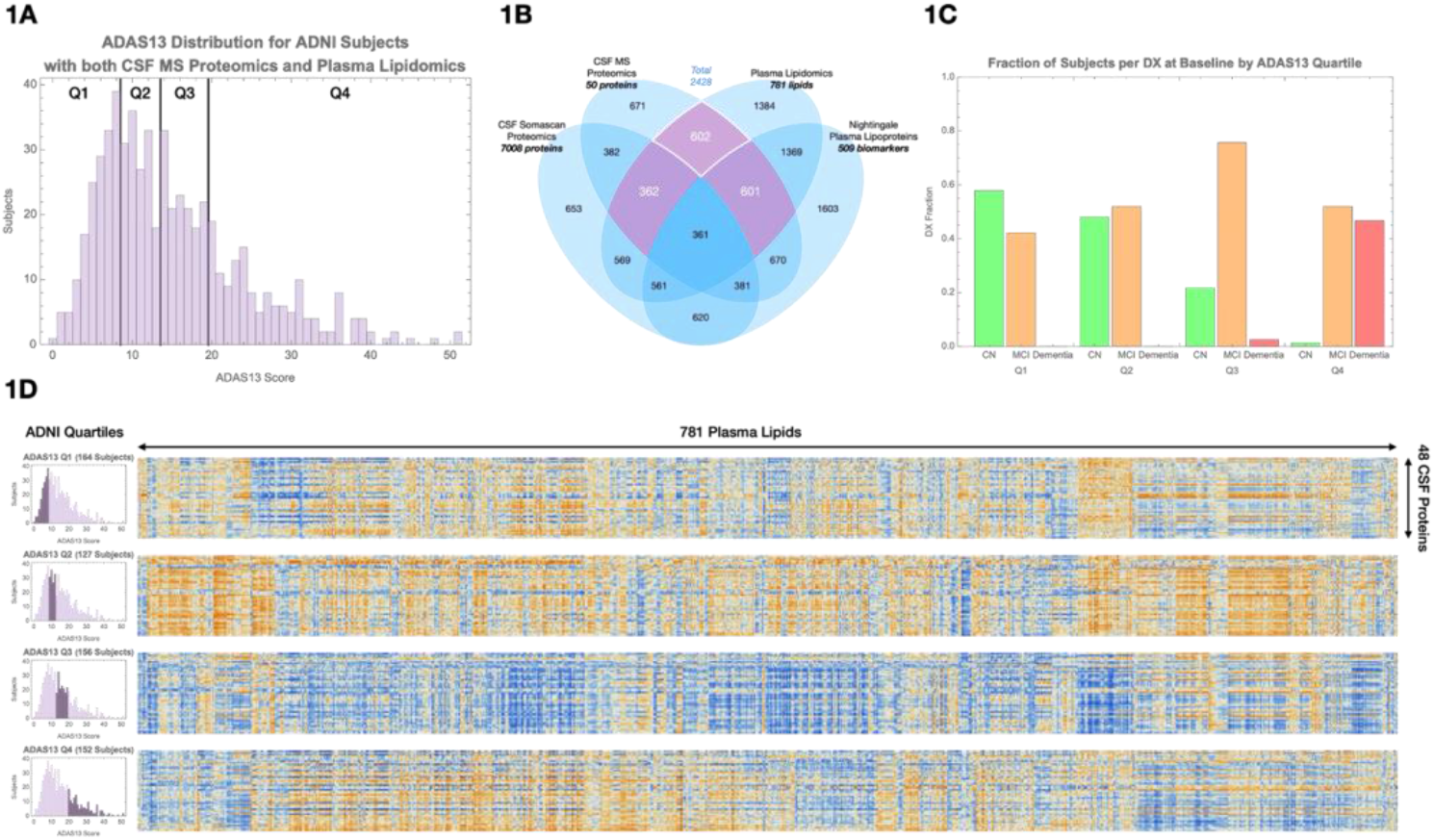
Overview of data selection and inter-omic correlation structure. **(A)** Schematic of ADNI multiomic datasets used in this study, including plasma lipidomics, CSF mass spectrometry (MS) proteomics, CSF SomaScan proteomics, and plasma lipoprotein profiles. **(B)** Selection of 602 participants with paired plasma lipidomic and CSF proteomic data; 362 had additional CSF SomaScan data, and 601 had Nightingale lipoprotein profiles. **(C)** Distribution of clinical diagnoses across cognitive quartiles based on ADAS13 scores: Q1 = 1–7, Q2 = 8–12, Q3 = 13–18, Q4 = 19–52. **(D)** Heatmaps of lipid–protein correlations within each cognitive quartile. Blue indicates negative correlations; orange indicates positive correlations. A visible shift in correlation patterns occurs between Q2 and Q3, corresponding to the cognitive transition from normal to mild impairment.

We selected 602 participants with paired baseline plasma lipidomic and CSF proteomic data^21^, including 362 participants with additional CSF SomaScan proteomic data^20^ and 601 with Nightingale plasma lipoprotein profiles (**Figure 1A–B**). Within this cohort, we analyzed correlations between 781 plasma lipids and 48 CSF proteins previously shown to differ between cognitively normal and AD participants^22–24^.

Correlation matrices were constructed separately within each cognitive quartile. Distinct shifts in lipid–protein associations emerged as cognitive scores declined, with a pronounced reversal in correlation sign (blue vs orange) between Q2 and Q3 corresponding to the cognitive transition from normal to MCI (**Figure 1D**).

To better organize these molecular changes, we performed unsupervised clustering of plasma lipids and CSF proteins. Plasma lipids grouped into ten clusters based on headgroup and fatty acid composition, including plasmalogens, cholesterol esters, and hexosylceramides (**Supplemental Table 1A**). CSF proteins grouped into three major clusters: complement/HDL-related proteins, hemoglobin-related proteins, and a larger cluster encompassing astrogliosis, ANLS/glycolysis, hypoxia, synaptic, and phagocytic markers.

Principal component analysis (PCA) was applied to each cluster to summarize major axes of variation across individuals. Correlations between the first principal components of lipid and protein clusters were evaluated within each quartile (**Figure 2A**), revealing specific disruptions in lipid metabolism and glial function as cognition declined.

**Figure 2.**
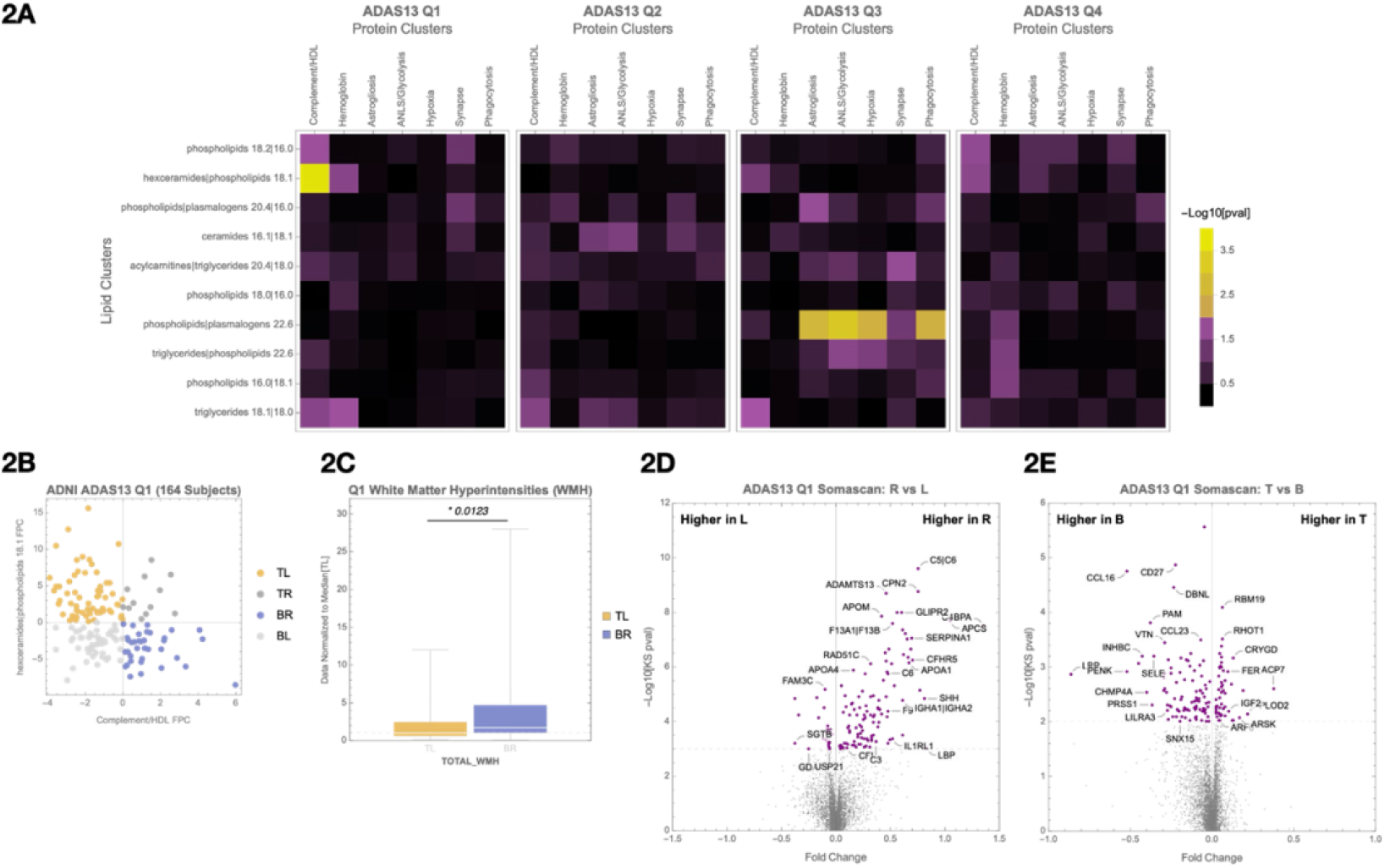
Cognitive quartile organization and principal component-based quadrant analysis of lipid and protein clusters. **(A)** Participants are grouped into cognitive quartiles based on ADAS13 scores. Within each quartile, correlations between selected plasma lipid and CSF protein clusters are evaluated using first principal component (FPC) values. **(B)** Scatterplot of participants’ lipid and protein FPC scores, illustrating quadrant definitions: top-left (TL), top-right (TR), bottom-left (BL), and bottom-right (BR). This quadrant structure is used for downstream molecular and clinical comparisons. **(C)** In Quartile 1 (Q1), comparison of white matter hyperintensity (WMH) burden between TL and BR quadrants. **(D)** In Q1, SomaScan differential expression analysis comparing Right vs Left quadrants (protein FPC axis). **(E)** In Q1, SomaScan differential expression analysis comparing Top vs Bottom quadrants (lipid FPC axis). Associated pathway enrichments are shown in **Supplementary Figures 2A (D) and 2B (E)**.

### Quartile 1 Analysis

In Q1, we observed early changes in the inter-omic correlation structure between plasma lipids and CSF proteins. A significant association (p = 1.2e-5) was identified between a lipid cluster enriched for hexosylceramides and phospholipids (“hexceramides | phospholipids 18.1”) and a CSF protein cluster composed of complement- and HDL-associated proteins, including ALB, APOA4, APOC1, APOC2, CP, C9, KNG1, PGLYRP2, PON1, and THRB (**Figure 2A, Q1**).

To visualize variation in these correlated signals, participants were divided into quadrants based on the first principal component (FPC) values of the lipid and protein clusters (**Figure 2B**). Comparing the bottom-right (BR) and top-left (TL) quadrants revealed a 58% higher white matter hyperintensity (WMH) burden in the BR group (**Figure 2C**), characterized by lower hexceramide and higher complement/HDL signals. WMH is a marker of vascular dysfunction and impaired BBB integrity, suggesting that shifts in lipid–protein coupling may precede overt cognitive symptoms^25,26^.

We next used SomaScan proteomics to compare molecular signatures between individuals in the right versus left quadrants of the plot (**Figure 2D**). This comparison revealed 72 differentially expressed CSF proteins (FDR < 0.05), with strong enrichment for complement and coagulation pathways (**Supplementary Figure 2A**). Pathway enrichment of the SomaScan proteins overlapped with those measured by mass spectrometry, suggesting that complement system activation is a convergent signal across platforms.

In a separate comparison of top versus bottom quadrants—defined by lipid cluster FPC—we identified 49 proteins (**Figure 2E, Supplemental Table 3A**). EnrichR identified the *Alternative Complement Pathway* as most significant (adj p = 3.3e-4) (**Supplemental Table 3B**).

### Quartile 3 Analysis

While Quartile 2 (Q2) showed some visible changes in overall inter-omic correlation structure (**Figure 1D**), no strong lipid–protein cluster correlations emerged when assessed by principal component analysis (**Figure 2A, Q2**). This suggests an early phase of systemic adjustments without system-wide disruption. In contrast, Quartile 3 (Q3) revealed striking and biologically coherent inter-omic correlation reversals, marking a major shift in homeostatic balance.

#### Astrogliosis protein cluster

We found a strong association in Q3 between the astrogliosis protein cluster (including YWHAB, YWHAZ, CHI3L1, and CD44) and the phospholipids | plasmalogens 22.6 lipid group (**Figure 3A**). CHI3L1 and CD44 are receptor–ligand partners linked to neuroinflammation and disease progression^27^, while YWHAB and YWHAZ bind GFAP, marking activated astrocytes^28^. Participants in the BR quadrant showed a 55% increase in brain TAU PET levels and a 92% increase in hippocampal atrophy rate compared to the TL quadrant (**Figure 3B**). These changes were accompanied by higher GFAP and GAS6 levels in the SomaScan data, known to be regulated by LXRA^29^.

**Figure 3.**
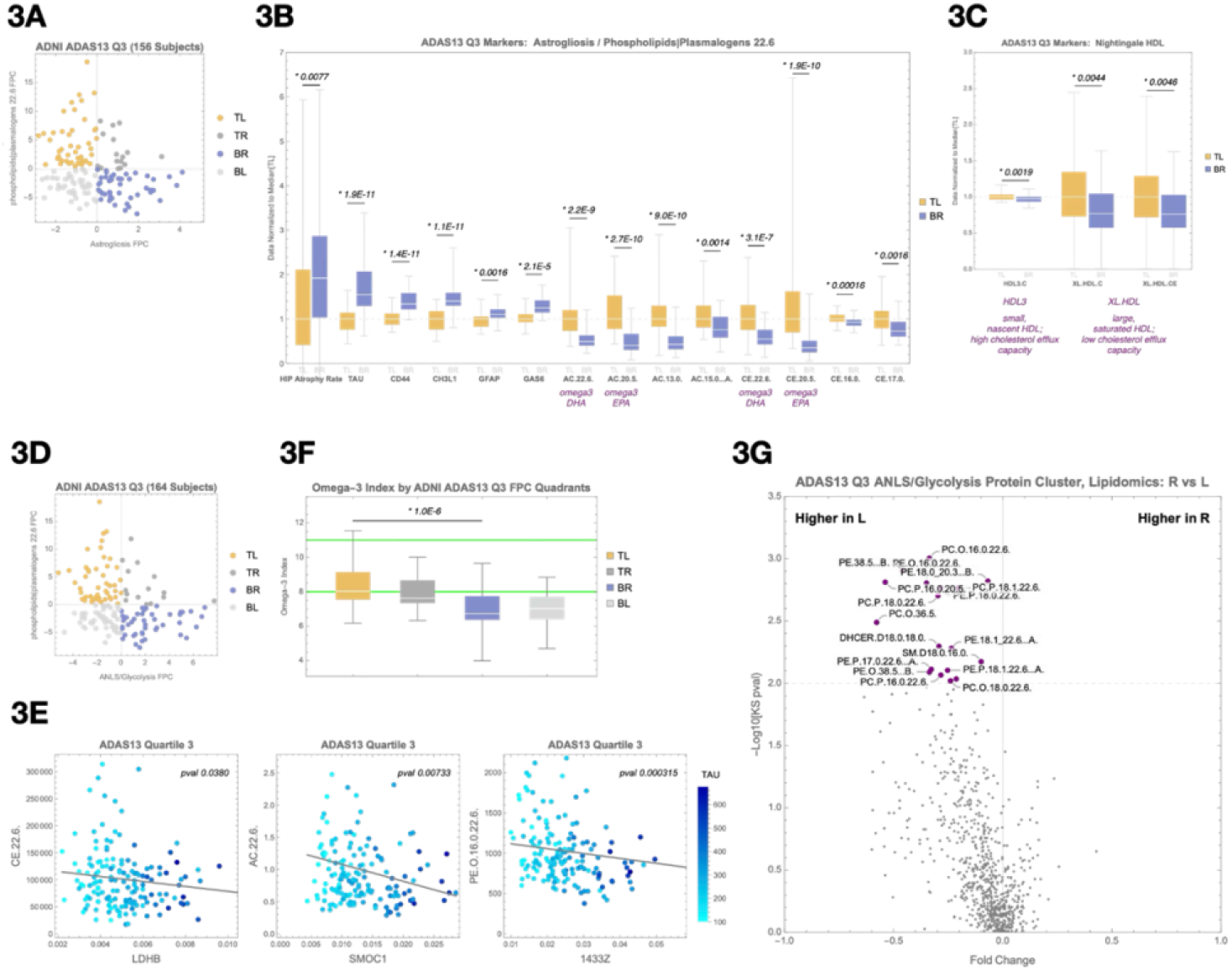
Emergence of lipid–protein associations linked to tau pathology and cerebrovascular changes in Quartile 3. **(A)** Principal component scatterplot of participants in Q3 based on the astrogliosis protein cluster (x-axis) and the phospholipids | plasmalogens 22.6 lipid cluster (y-axis). **(B)** Box-and-whisker plots comparing the top-left (TL) and bottom-right (BR) quadrants from (A) across markers of disease progression (hippocampal atrophy rate, tau PET signal), astrogliosis (CD44, CHI3L1, GFAP), beta oxidation (omega-3 and non–omega-3 acylcarnitines), and reverse cholesterol transport (omega-3 and non– omega-3 cholesterol esters). **(C)** Box-and-whisker plots comparing TL and BR quadrants from Nightingale NMR HDL lipoprotein markers, including HDL3, XL.HDL.C, and XL.HDL.CE. **(D)** Principal component scatterplot of participants based on the ANLS/glycolysis protein cluster (x-axis) and the phospholipids | plasmalogens 22.6 lipid cluster (y-axis). **(E)** Representative protein–lipid correlations in Q3, including LDHB–CE.22.6, SMOC1–AC.22.6, and YWHAB (14-3-3 zeta)–PE(O-16:0/22:6) associations. These examples illustrate disrupted coupling between lipid metabolism and astrocyte or glycolytic protein markers under metabolic stress. **(F)** Comparison of the estimated omega-3 index, calculated from DHA- and non-DHA–containing HDL particles measured by Nightingale NMR, between BR and TL quadrants. Median omega-3 index was significantly lower in the BR quadrant (p = 8.6e-7), suggesting impaired omega-3 availability may contribute to disrupted plasmalogen biosynthesis and astrocyte–neuron metabolic support. **(G)** SomaScan differential expression analysis comparing Right vs. Left quadrants from (A), visualized as a volcano plot. Notable proteins enriched in the BR quadrant include GFAP and GAS6, consistent with astrocyte activation.

Supporting impaired lipid metabolism, the BR quadrant also showed increased acylcarnitines and cholesterol esters, suggesting reduced lipid oxidation and cholesterol efflux (**Figure 3B,C**). To determine whether these elevated lipid species could be traced to the brain, we incorporated plasma lipoprotein data from the Nightingale NMR platform. Notably, the TL quadrant showed higher levels of a CNS-specific extremely large HDL particle — a lipoprotein particle class known to originate exclusively from the brain^30,31^. The detection of this CNS-derived particle in peripheral blood provides key evidence that lipid signals observed in plasma may reflect active export from the brain. This observation supports the idea that microglial lipid processing capacity is exceeded, with reduced cholesterol and lipids trafficked out of the CNS into circulation.

#### ANLS/glycolysis protein cluster

The strongest protein–lipid association in Q3 was between the ANLS/glycolysis protein cluster (including LDHB, LDHC, CD44, ALDOA, PKM, MDH1, GAA, and ENO1) and the lipid cluster phospholipids | plasmalogens 22.6, which includes many Docosahexaenoic acid (DHA)- and Eicosapentaenoic acid (EPA)-containing lipids (**Figure 3D**). Several individual protein–lipid pairs within this quadrant showed significant associations, including LDHB and CE.22.6 (**Figure 3E**). Participants in the BR quadrant, characterized by higher glycolysis and lower plasmalogen signals, exhibited a 65% higher median brain TAU PET level than those in the TL quadrant (**Supplementary Figure 3**). These results support the hypothesis that impaired lipid processing results in metabolic stress as reflected in compensatory changes in glycolytic proteins.

To probe whether this metabolic vulnerability was related to omega-3 availability, we estimated the omega-3 index using Nightingale NMR data on DHA- and non-DHA–containing HDL particles^32–34^. The median omega-3 index in the TL quadrant was within the clinically recommended range, while the BR quadrant had a significantly lower median (p = 8.6e-7) (**Figure 3F**). This finding aligns with the known protective effects of omega-3s against AD progression and highlights a mechanistic link: omega-3s and plasmalogens are critical for lipid droplet trafficking^35,36^.

Taken together, these data reveal a coherent axis connecting decreased omega-3 and plasmalogen levels with disrupted expression of ANLS and glycolytic proteins. This dysfunction appears to originate from a buildup of neuronal membrane debris—including myelin fragments—that must be cleared by glia. When the burden of this cholesterol-rich material exceeds the trafficking capacity of astrocytic lipoproteins and lipid droplet systems, astrocytes become overloaded. The resulting membrane saturation impairs essential functions such as GPCR signaling and cholesterol export, disrupting astrocyte–neuron communication. In particular, ANLS becomes compromised, depriving neurons of a key metabolic support pathway.

This metabolic fragility may represent a critical inflection point that triggers early tau accumulation and neurodegeneration. Prior studies show that neurons under energetic stress activate AMP-activated protein kinase (AMPK) to compensate for ATP deficits^37–39^. However, in the absence of sufficient astrocytic support, AMPK activation becomes maladaptive— contributing to calcium dysregulation and tau phosphorylation^40–42^. This cascade promotes synaptic dysfunction and accelerates neurodegenerative progression.

### Quartile 2 Analysis

The strong astrocytic and tau pathology signals in Quartile 3 (Q3) prompted us to revisit Quartile 2 (Q2) to search for earlier signs of metabolic stress and impaired cholesterol flux — prior to measurable cognitive decline or glial activation. Although no significant lipid–protein cluster correlations emerged at the FPC level (**Figure 2A, Q2**), we identified several FPC pairs with low p-values and investigated their component lipids and proteins for meaningful associations.

This analysis revealed a consistent association between the ANLS/glycolysis protein cluster and a ceramide-rich lipid cluster, specifically centered around CER.D19.1.18.0, an odd-chain sphingolipid. We stratified participants by principal component scores from these two clusters (**Figure 4A**), revealing a 59% increase in brain TAU PET and higher hippocampal atrophy rate in the bottom-right (BR) quadrant compared to the top-left (TL) (**Figure 4B**). Among individual correlations, LDHB–CER.D19.1.18.0 was the strongest (p = 0.0046), with broad correlations seen across glycolytic enzymes (**Figure 4C**).

**Figure 4.**
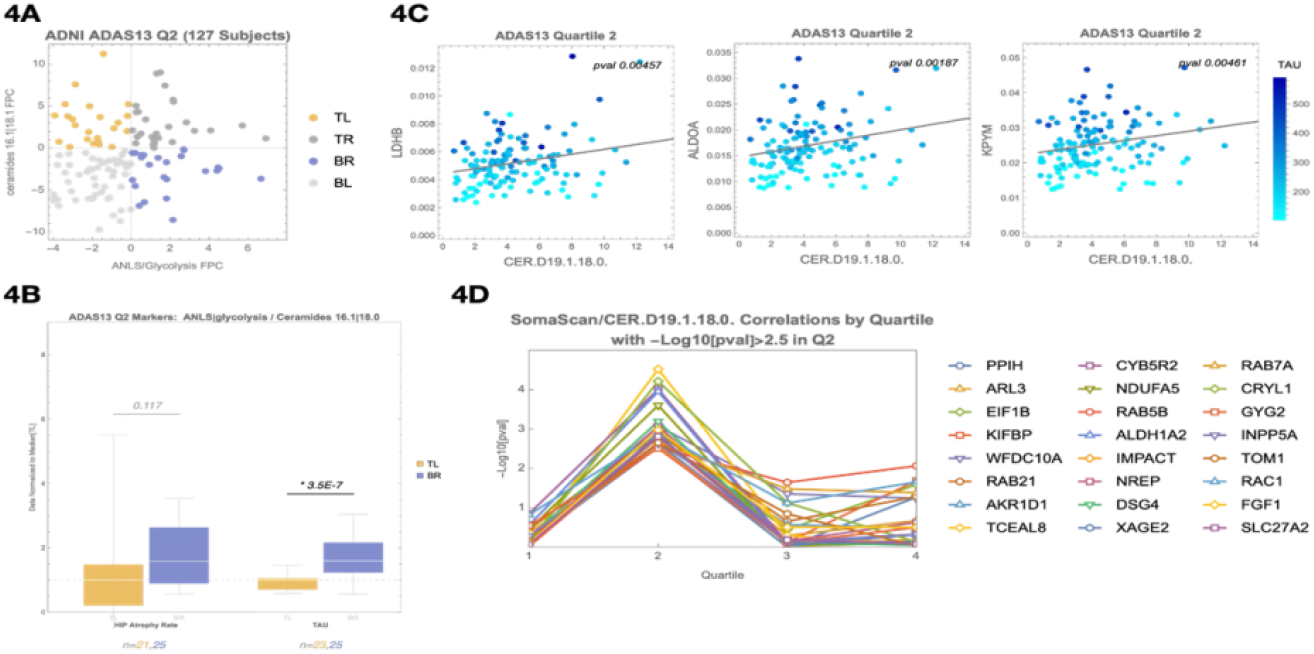
A ceramide-enriched lipid cluster, including CER.D19, emerges in Quartile 2 as a marker of flux imbalance and disrupted astrocyte–neuron metabolic support. **(A)** PCA-based quadrant analysis of the ANLS/glycolysis protein cluster (x-axis) and a ceramide-enriched lipid cluster (y-axis) in Q2. Participants in the bottom-right (BR) quadrant exhibit higher ANLS/glycolysis protein expression and lower ceramide lipid signal, suggesting a mismatch between increasing metabolic demand and insufficient lipid clearance. **(B)** Comparison of tau PET signal and hippocampal atrophy rate between TL and BR quadrants defined in (A). Participants in the BR quadrant show a 59% increase in tau PET signal and elevated atrophy rate. **(C)** Scatter plots showing correlations between CER.D19.1.18.0 and representative ANLS/glycolysis proteins (LDHB, ALDOA, KPYM) in Q2. LDHB–CER.D19.1.18.0 is the strongest association (p = 0.0046), supporting disrupted metabolic coupling at this stage. **(D)** SomaScan differential expression analysis of CSF proteins associated with CER.D19.1.18.0 levels. Top proteins include RAB21, RAB7A, SLC27A2, RAC1, and FGF1 — involved in lipid droplet trafficking, cholesterol esterification, and oxysterol signaling. These associations were specific to Q2.

To explore the biology behind CER.D19, we performed a proteomic correlation screen using SomaScan CSF data. We identified 24 proteins significantly associated with CER.D19.1.18.0 (p < 0.003), many of which participate in lipid droplet trafficking (RAB21, RAB7A)^43^, cholesterol esterification (SLC27A2)^44–46^, and oxysterol signaling (RAC1, FGF1, INPP5A)^47^. These were notably Q2-specific and not observed in Q1 or Q3 (**Figure 4D; Supplemental Table 4A**). Pathway analysis via EnrichR identified bile acid biosynthesis via 24-hydroxycholesterol as the top hit (adjusted p = 6.1e-3; **Supplemental Table 4B**). The CER.D19 odd-chain fatty acid is synthesized using propionyl-CoA, a byproduct of cholesterol side-chain β-oxidation during bile acid synthesis^48–56^.

This is significant because the conversion of cholesterol to 27-hydroxycholesterol represents one of the routes for cholesterol export from the brain, enabling it to cross the blood– brain barrier and be metabolized into bile acids in the liver. The proteins identified here map directly to the molecular machinery required for this process, suggesting that CER.D19 levels correlate with the point where glial cells begin activating compensatory mechanisms to manage excess cholesterol through this export route.

Given this connection to bile acid metabolism, we asked whether any known AD-associated risk loci might be enriched among individuals with altered CER.D19 levels. To do this, we curated a list of 101 genome-wide significant AD GWAS loci from recent meta-analyses and tested for associations with CER.D19.1.18.0. This approach revealed variants near PICALM, INPP5D, TSPOAP1, and TNIP1—all genes implicated in cholesterol trafficking, mitochondrial lipid processing, or bile acid synthesis^57^ (**Figure 5C; Supplemental Figure 4A**). Additionally, SNP rs686548 in SPTLC3, the rate-limiting enzyme for odd-chain ceramide synthesis, was identified as the top CER.D19-associated SNP via Baker PheWeb (**Supplemental Figure 4B**). Because of the known role of LXRA in lipid metabolism in the mitochondria^38^, we also looked to see if any LXRA-associated (NR1H3) SNPs correlated and found rs7120118 and rs3758673 to correlate with CER.D19.1.18.0.

**Figure 5.**
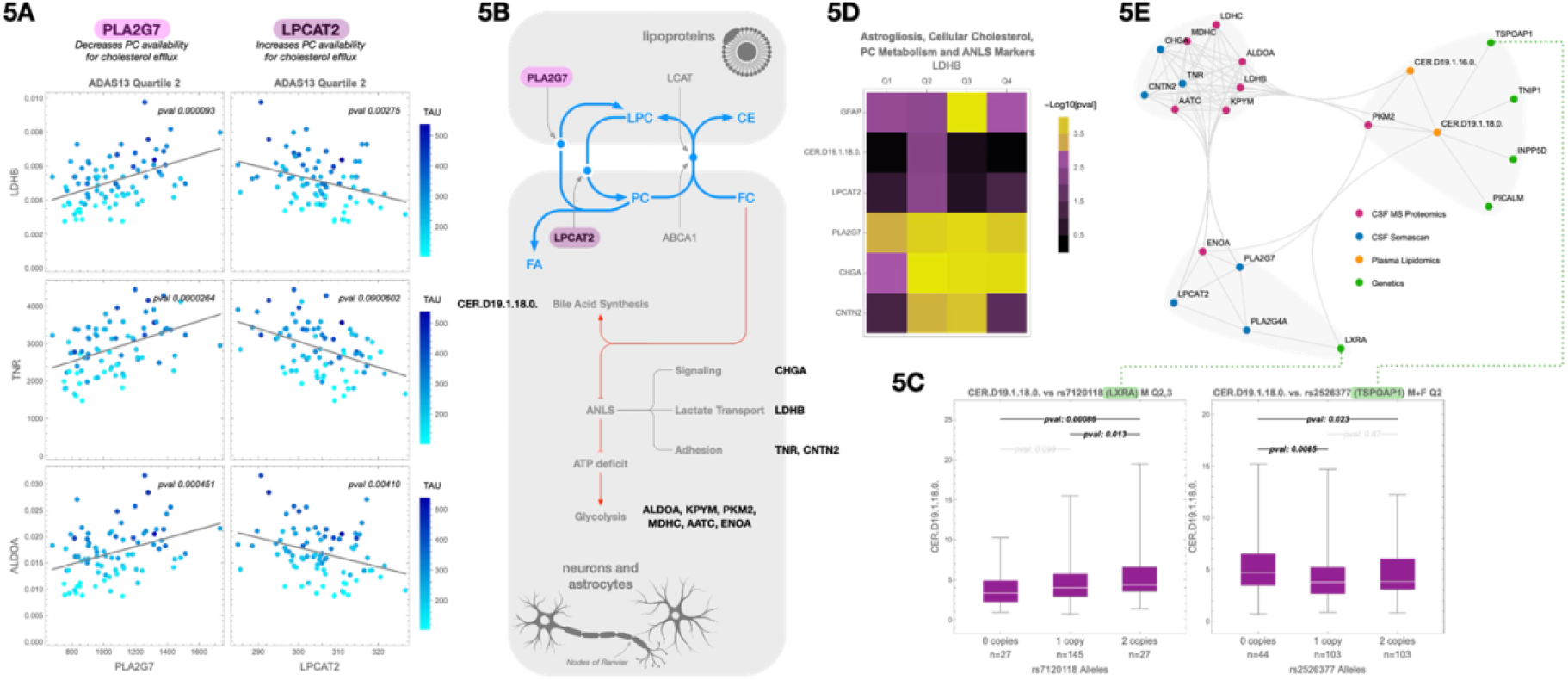
CER.D19 marks a metabolic inflection point linked to cholesterol overload, bile acid biosynthesis, and GWAS-enriched lipid handling pathways. **(A)** Correlations between CSF LDHB (a marker of astrocyte–neuron lactate shuttle activity), TNA and ALDOA and two enzymes involved in cholesterol esterification: PLA2G7 (positive correlation) and LPCAT2 (negative correlation). These associations suggest altered phosphatidylcholine (PC) processing and potential constraints on cholesterol export. **(B)** Schematic illustration showing how PLA2G7 and LPCAT2 regulate the interconversion of PC and lysophosphatidylcholine (LPC), affecting cholesterol solubilization and packaging into APOE-containing lipoprotein particles for efflux. **(C)** Representative AD GWAS SNPs associated with CER.D19.1.18.0 levels, including loci near NR1H3 (LXRA) and TSPOAP1, which regulate nuclear cholesterol sensing and mitochondrial cholesterol trafficking, respectively. Additional associated loci are provided in **Supplemental Figure 4A and Supplemental Table 4C**. **(D)** Heatmap showing correlations between CSF LDHB and selected markers across all cognitive quartiles. Variables include GFAP (astrogliosis), CER.D19.1.18.0 (cellular cholesterol load), PLA2G7 and LPCAT2 (PC metabolism), and CHGA and CNTN2 (ANLS signaling and adhesion). **(E)** Summary diagram integrating lipidomic, proteomic, and genetic findings: early CER.D19 elevation coincides with metabolic stress markers (e.g., ANLS disruption), cholesterol overload, bile acid biosynthesis, and the emergence of AD risk gene expression signatures consistent with declining microglial lipid flux capacity.

These genetic findings reinforce the proteomic evidence, pointing to upstream regulatory steps in cholesterol clearance. Many of these genes control lipid droplet formation, vesicular trafficking, and mitochondrial cholesterol import—critical steps that precede conversion of cholesterol to 27-hydroxycholesterol for LXRA and SREBP2 signaling^11,12^. Their enrichment among individuals with altered CER.D19 levels suggests a genetically influenced bottleneck in cholesterol flux capacity that may predispose certain individuals to early metabolic breakdown in AD.

We next examined cholesterol handling enzymes and their relationship to ANLS/glycolysis proteins. LDHB, a central member of this protein cluster, serves as an indicator of neuronal metabolic stress and astrocyte–neuron metabolic coupling^58,59^. Notably, LDHB showed a positive correlation with CSF PLA2G7 and a negative correlation with LPCAT2 (**Figure 5A**), consistent with the opposing roles these enzymes play in phosphatidylcholine (PC) metabolism and cholesterol esterification^60,61^. These relationships suggest that dysregulated PC turnover—via increased PLA2G7 or decreased LPCAT2—could impair cholesterol packaging into APOE lipoproteins, leading to astrocytic overload and downstream metabolic stress.

Additional CSF markers associated with CER.D19 included CNTN2 and CHGA, both of which also correlated with LDHB across quartiles (**Figure 5D**). CNTN2, a cell adhesion protein that localizes to nodes of Ranvier, peaked in Q2 and Q3, while CHGA, co-secreted with norepinephrine, remained stable from Q2 through Q4^62^. These proteins are involved in astrocyte–neuron communication and reflect the integrity of GPCR-mediated signaling cascades. LDHB, a key member of the ANLS/glycolysis protein cluster, serves as an indicator of neuronal metabolic stress and astrocyte–neuron metabolic decoupling^38^. Importantly, excess cholesterol in astrocytic membranes has been shown to impair GPCR signaling^63,64^, including pathways regulated by norepinephrine and glutamate, which are critical for triggering ANLS^65^.

Thus, the co-variation of LDHB with CNTN2 and CHGA suggests that rising intracellular cholesterol may begin to impair signaling at precisely the point when microglial cholesterol processing begins to saturate. A summary of these integrated lipidomic, proteomic, and genomic correlations is shown in **Figure 5E**.

Taken together, these results suggest that Q2 is not a silent interval, but a biologically active phase of incipient flux imbalance, in which microglia lipid handling begins to saturate. CER.D19 appears to mark this metabolic tension point, linking excess intracellular cholesterol with early disruption of signaling-based astrocyte–neuron coordination and compensatory mitochondrial processing. This lipid accumulation appears to interfere with GPCR signaling necessary for lactate shuttling via the ANLS, leading to rising LDHB levels as neurons experience mounting metabolic stress. These changes precede overt glial activation and tau pathology, reinforcing the hypothesis that early cholesterol buildup—not depletion—initiates the breakdown of homeostatic support systems in AD.

## Methods

### Data availability statement

The results published here are in whole or in part based on data obtained from the AD Knowledge Portal (https://adknowledgeportal.org).

ADNI metabolomics data from the Baker lipidomics kit is available at the AD Knowledge Portal under https://adknowledgeportal.synapse.org/Explore/Studies/DetailsPage?Study=syn5592519 and https://adknowledgeportal.synapse.org/Explore/Studies/DetailsPage/StudyDetails?Study=syn24989039, the full complement of clinical and demographic data for the ADNI cohorts are hosted on the LONI data sharing platform and can be requested at http://adni.loni.usc.edu/data-samples/access-data/.

### ADNI Data

All ADNI data was downloaded from https://adni.loni.usc.edu/

### ADNI Genetics

A harmonized array dataset for ADNI has been published previously^30,31^. Briefly, genotype array data from ADNI1, ADNI2/GO, ADNI Omni 2.5, and ADNI 3 were processed and imputed using the TOPMED imputation server. Variants were removed for missingness (5%), low minor allele frequency (<1%), or if out of Hardy Weinberg Equilibrium. Samples were removed for missingness (1%), relatedness (pi-hat>.25), inconsistency between reported sex and genetic sex, and for heterozygosity (>5standard deviations). Imputed data were called for variants with high imputation quality (R2>0.80). Participants that were present in more than one array were dropped from the array of lower quality. After imputation and QC, 1,559 participants and 8,028,923 variants were available for analysis.

### Baker Metabolomic PhWAS

https://metabolomics.baker.edu.au/pheweb_standard/region/Cerd191160/20:12773521-13173521

### Statistical analysis

Code for all the accompanying analysis can be found in the accompanying Mathematica files

### Clustering and Principal Component Analysis

Clustering, principal component analysis, linear regressions, and statistical analyses were performed using Wolfram Mathematica v14.0.

### Hippocampal Atrophy Rate Calculation

For each individual, hippocampal atrophy rate was calculated by linear regression over all available time points measures of hippocampus volume, and presented as percent per year normalized to volume at baseline.

### Omega-3 Index Calculation

We calculated the omega-3 index based on the description found in Schuchardt and et al Schacky et al^66^. Briefly, we log-transformed the O3I and/or DHA% and non-DHA% prior to fitting a regression equation. The additive model containing two terms was determined to be the final prediction model, including β-coefficients for term: eO3I = 2·629 × NMR DHA% þ 0·4673 × NMR non-DHA% − 0·1014. The correlation of the eO3I vs observed O3I values was calculated for all ADNI participants.

## Discussion

AD has long been conceptualized as a proteinopathy, molecularly defined by extracellular amyloid-beta plaques and intracellular tau tangles. While these hallmark pathologies are correlated with disease progression and cognitive decline, there remains a disconnect between their presence and our mechanistic understanding of disease etiology. This has led to growing interest in metabolic and glial contributions to early vulnerability. In this study, we applied integrated lipidomic and proteomic profiling of CSF and matched plasma samples— stratified by cognitive performance quartiles—to uncover early molecular changes consistent with impaired cholesterol efflux. These were accompanied by genetic associations and proteomic shifts implicating microglial lipid handling, suggesting that a failure in metabolic flux capacity precedes and potentiates the downstream cascades commonly associated with AD.

Our findings converge on a model of progressive collapse in cholesterol and lipid homeostasis, driven by overwhelmed glial flux capacity. In Quartile 1 (Q1), lipid–protein correlation structures were still largely intact, with subtle shifts suggesting early adaptive changes. Notably, several of the protein and lipid clusters showing early correlation shifts are enriched for markers of vascular integrity and hemoglobin metabolism, consistent with prior reports of early BBB disruption in AD^47^. These associations suggest that vascular leakage may precede metabolic strain and may reflect compensatory responses to reduced cerebral perfusion.

In Quartile 2 (Q2), the absence of system-wide lipid–protein correlations initially suggested a transitional or uninformative phase. However, retrospective analysis motivated by the strong astroglial and metabolic signals observed in Q3 revealed that Q2 is in fact a critical dual inflection point. The emergence of CER.D19, an odd-chain ceramide linked to propionyl-CoA metabolism^48–56^, cholesterol esterification, and mitochondrial cholesterol trafficking, marked the beginning of detectable metabolic stress. Participants with elevated glycolytic protein expression and elevated CER.D19 levels showed higher tau burden and hippocampal atrophy, suggesting that cholesterol clearance capacity was disrupting neuronal metabolic support.

These observations were reinforced by the appearance of CNS-specific HDL markers in plasma and correlations between CER.D19 and proteins involved in bile acid biosynthesis and oxysterol metabolism. Furthermore, GWAS loci associated with CER.D19 levels included variants near several established AD risk genes: TSPOAP1, INPP5D, PICALM, TNIP1, and both NR1H3 (LXRA), and SPTLC3. These genes are involved in mitochondrial cholesterol trafficking, vesicular transport, nuclear lipid sensing, lipid droplet processing, and includes enzymes that produce CER.D19 as a byproduct of broader lipid metabolic processes^67^. Their enrichment supports the hypothesis that disrupted cholesterol handling is a genetically encoded vulnerability in AD and, more significantly, provides functional context for how these GWAS loci contribute to disease etiology—something that has remained elusive for many AD risk variants. Together, these data suggest that CER.D19 acts as a sentinel lipid, reflecting the point at which glial lipid flux systems begin to saturate.

By Quartile 3 (Q3), compensatory mechanisms showed widespread changes, with clear signs of metabolic and glial strain. Plasma cholesterol ester levels were inversely correlated with astrocyte-derived CNS protein markers, and participants with lower cholesterol ester signal and higher astrocytic stress showed increased tau burden. These patterns suggest that astrocytes are assisting with processing of excess cell debris, buffer excess lipids, impairing their ability to provide metabolic support for neurons via ANLS.

Our model shifts the focus of early AD pathogenesis away from amyloid and toward the failure of glial lipid processing and cholesterol clearance. Importantly, our results highlight a mechanistic explanation for the long-observed protective effects of omega-3 fatty acids: participants with higher omega-3 index showed greater preservation of plasmalogens, improved HDL profiles, and reduced tau pathology^68^. These effects likely stem from enhanced lipid droplet trafficking and membrane remodeling. Omega-3 polyunsaturated fatty acids, particularly DHA, contain multiple cis double bonds that introduce curvature into the lipid structure, promoting the formation of smaller, more dynamic lipid droplets^69^. This biophysical property facilitates mitochondrial docking and efficient cholesterol oxidation via oxysterol synthesis, thereby enhancing cholesterol efflux. In this way, omega-3s not only improve membrane composition but also mitigate cholesterol overload in astrocytes, helping preserve metabolic support to neurons during early stage disease.

Our unifying hypothesis posits that AD arises when the brain’s capacity to process, store, and export cholesterol is overwhelmed. Initially, astrocytes and microglia buffer this excess through lipid droplet formation and metabolism. However, as cholesterol-rich neuronal debris accumulates—particularly from aging and myelin turnover—these clearance systems become saturated. Enlarged and persistent lipid droplets disrupt their dynamic interactions with mitochondria, impairing lipid transfer and mitochondrial function. This breakdown leads to mitochondrial stress, disrupted ANLS, and subsequent tau aggregation. Rather than viewing amyloid and tau as primary instigators, we propose they are downstream consequences of unresolved lipid accumulation and failed clearance mechanisms.

Our hypothesis also provides a coherent framework to reinterpret longstanding genetic observations. APOE4, the strongest genetic risk factor for late-onset AD, has a shorter half-life^70^ and reduced lipid-carrying capacity^71^ compared to other isoforms, as well as altered binding affinities for LDLR and LRP1^71^. These differences impair the recycling of cholesterol-rich APOE particles back into astrocytes, increasing the likelihood of astrocytic cholesterol overload and impairing ANLS. In contrast, APOE2 carriers show more efficient lipid recycling and reduced astrocytic burden, preserving metabolic support, likely due to their reduced affinities for LDLR and LRP1^72^. The Christchurch mutation (APOE3 R136S), which disrupts APOE binding to heparan sulfate proteoglycans, likely reduces retention of cholesterol-laden particles in the extracellular space and inhibits their uptake, protecting astrocytes from saturation. This mechanism offers a plausible explanation for why the Christchurch variant mitigates the impact of a pathogenic PSEN1 mutation^11,73–75^—by preserving cholesterol flux equilibrium and delaying the onset of glial and neuronal dysfunction despite high amyloid burden.

These genetic effects align with our data, where rising CER.D19 levels are not interpreted as causal drivers of pathology, but rather as sentinel markers of disrupted cholesterol metabolism and flux capacity. Importantly, our findings intersect with known regulators of cholesterol homeostasis, including LXRA and SREBP2^11^. Under conditions of excess lipid accumulation, particularly when lipid droplets become enlarged and fail to fuse with mitochondria, mitochondrial β-oxidation is impaired. This disruption prevents 27-OHC synthesis and feedback inhibition of SREBP2, allowing de novo cholesterol synthesis to continue unchecked—even in the setting of cholesterol overload. Julia TCW and colleagues previously demonstrated that impaired lipid droplet turnover maintains active SREBP2 signaling and sustained cholesterol production^33,68^. Omega-3 fatty acids may counteract this process by preventing the formation of oversized lipid droplets, thereby maintaining droplet–mitochondrial interactions and allowing appropriate downregulation of SREBP2^31,76,77^. This mechanism helps preserve cholesterol homeostasis and may underlie the protective metabolic effects associated with higher omega-3 index^78^.

Our unified hypothesis integrates these findings into a model of Alzheimer’s pathophysiology, outlined in Figure 6. The process begins when microglia phagocytose neuronal and myelin debris, loading debris-derived cholesterol onto de novo-synthesized APOE lipoproteins (Step 1). These lipoproteins enter a shared pool with astrocyte-synthesized particles, but as particle size increases due to debris loading, cholesterol efflux capacity declines (Step 2). The accumulation of cholesterol in astrocytes and neurons impairs membrane dynamics, disrupts GPCR signaling, and reduces ANLS-mediated lactate delivery to neurons (Step 3). Neurons experiencing metabolic stress activate AMPK but cannot compensate for the energetic shortfall^39,79^, leading to calcium dysregulation^79,80^, tau phosphorylation, and progressive neurodegeneration (Step 4). As microglial flux capacity saturates, astrocytes assume a phagocytic role, accelerating cholesterol accumulation and triggering astrogliosis (Step 5). The final bottleneck occurs when cholesterol processing stalls at the level of lipid droplet–mitochondrial coupling, a step genetically influenced by loci such as INPP5D, TSPOAP1, PICALM, TNIP1, SPTLC3, and NR1H3 (LXRA), all of which correlated with CER.D19.1.18.0. levels in our analysis (Step 6).

**Figure 6.**
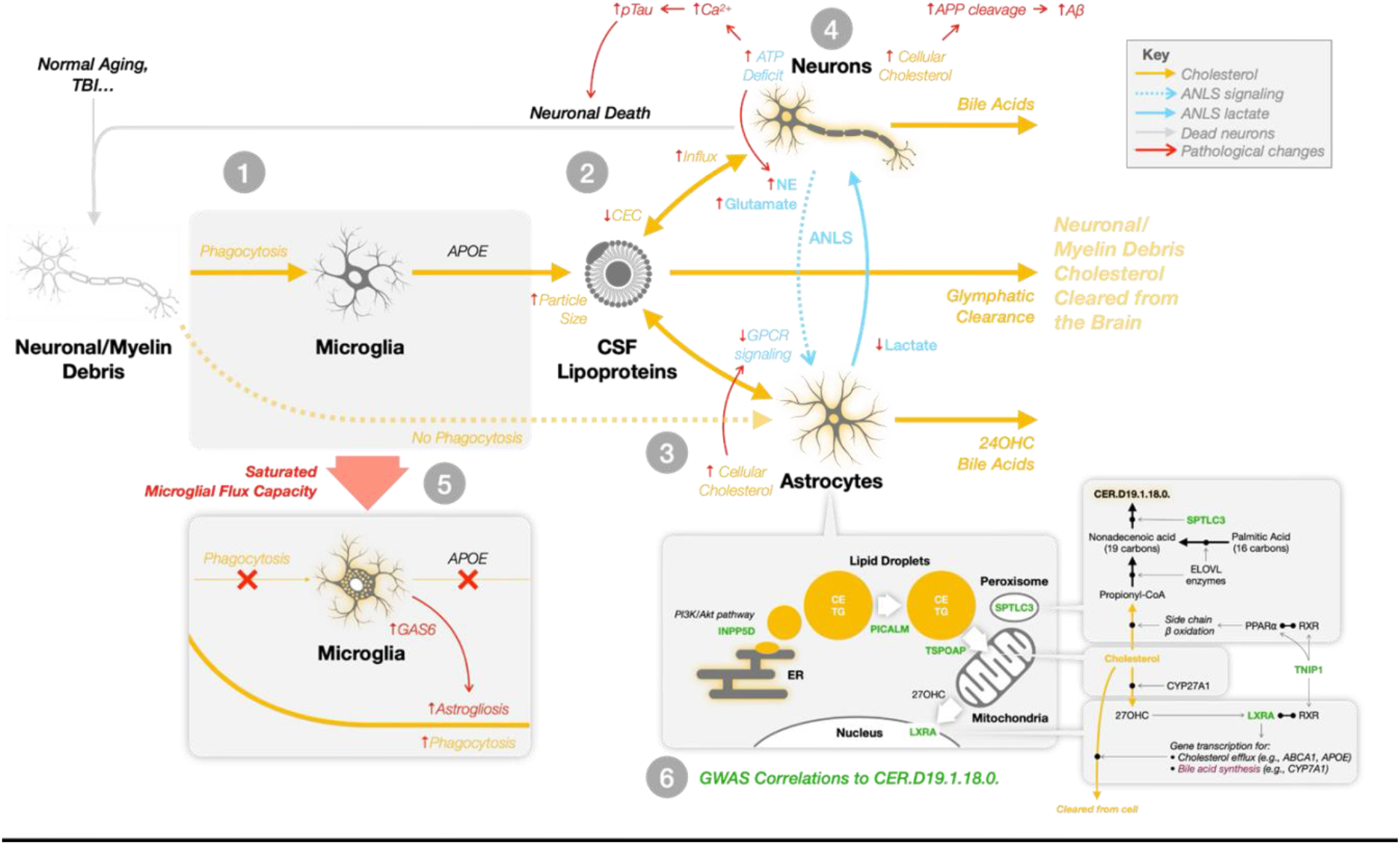
Unified hypothesis of Alzheimer’s disease: progressive collapse of microglial cholesterol flux capacity and astrocyte-neuron metabolic breakdown. This schematic outlines a multi-step systems-level failure in brain lipid homeostasis that we propose underlies Alzheimer’s disease progression. **1:** Microglia phagocytose cholesterol-rich debris from neurons and myelin. This cholesterol is packaged into newly synthesized APOE lipoproteins. **2:** These debris-laden particles merge with the existing pool of astrocyte-derived APOE lipoproteins, which are responsible for transporting astrocyte-synthesized cholesterol to neurons. As the size of lipoproteins increases due to debris accumulation, their cholesterol efflux capacity (CEC) via ABCA1 decreas limiting the removal of excess neuronal cholesterol. **3:** Elevated cholesterol levels in neurons and astrocytes impair membrane dynamics. In neurons, this drives APP cleavage to amyloid-beta; in astrocytes, cholesterol buildup disrupts GPCR signaling, impairing the astrocyte–neuron lactate shuttle (ANLS) and reducing neuronal access to metabolic fuel. **4:** Metabolically vulnerable neurons attempt to compensate through AMPK activation, but remain under-energized. Resultant calcium dysregulation leads to hyperphosphorylation of tau and other stress responses, promoting a feedforward loop of neurodegeneration. **5:** As microglial flux capacity is exhausted, astrocytes assume direct phagocytic roles. This induces astrogliosis and further elevates astrocytic cholesterol burde amplifying metabolic disruption and inflammatory signaling. **6:** Genetic variants at loci including *INPP5D, TSPOAP1, PICALM, TNIP1, NR1H3* (LXRA), and *SPTLC3* affect key steps in intracellular cholesterol trafficking and lipid droplet processing. These processes determine whether cholesterol can be shuttled to mitochondria for oxidation and bile acid synthesis or instead accumulates pathologically. Many of these genes were significantly correlated with CER.D19.1.18.0, an odd-chain ceramide that serves as a marker of flux bottleneck and mitochondrial metabolic failure. This model illustrates how failure to clear excess cholesterol disrupts both structural and signaling roles of membrane lipids, deprives neurons of metabolic

Our model reframes AD as a failure of coordinated lipid handling and metabolic support, upstream of amyloid and tau deposition. It provides a mechanistic framework to explain decades of AD genetics, proteomics, and lipidomic findings, while unifying disparate observations into a coherent system-level pathology. The root cause of vulnerability may lie not in protein aggregation per se, but in the brain’s failure to remove cholesterol debris without dysregulating homeostatic processes. The appearance of CER.D19 and associated proteomic and genomic signals in Q2 provides a mechanistic entry point into this process. This research emerged from our efforts to construct digital twins of aging individuals. While our models performed well in predicting biomarker trajectories during normal cognition, their accuracy deteriorated upon reaching the mild cognitive impairment (MCI) phase. Our digital twin models consistently pointed to unexplained variability associated with cholesterol metabolism and lipid handling, particularly in glial compartments. This led us to hypothesize that a key missing axis of variation was microglial flux capacity—the ability of glial cells to manage cholesterol clearance and maintain metabolic equilibrium. The findings presented here validate that hypothesis and are now being integrated back into our digital twin framework to improve forecasting of disease progression and better capture the early transition points preceding overt pathology, work that will be presented separately due to its distinct scope and intention.

Taken together, these findings suggest that AD may benefit from being reframed in terms of etiology—as a systems-level breakdown in cholesterol flux and glial metabolic capacity, rather than solely a disorder of protein aggregation. Our hypothesis offers a potentially unifying framework through which many known risk and resilience factors—including genetic variants, lipid signatures, and environmental modifiers—can be interpreted. While it does not resolve the long-standing debate over AD causation or definition, it contributes a mechanistically grounded perspective focused on the biophysical dynamics of cholesterol. By incorporating concepts of mass balance, metabolic flux, and feedback regulation, this model helps contextualize the complex and heterogeneous molecular patterns observed across patients, and can guide future efforts to refine our understanding of AD etiology, progression and personalized therapeutic strategies.

## Abbreviations

AD: Alzheimer’s disease
ADNI: Alzheimer’s Disease Neuroimaging Initiative
GWAS: genome wide association study ANLS astrocyte-neuron lactate shuttle
LD: lipid droplets
24-OHC: 24-hydroxycholesterol
27-OHC: 27-hydroxycholesterol
CSF: cerebrospinal fluid
HDL: high density lipoprotein
LDL: low density lipoprotein
WMH: white matter hyperintensity
ADNI: Alzheimer’s Disease Neuroimaging Initiative
ADAS13: Alzheimer’s Disease Assessment Scale-Cognitive 13
PCA: principle component analysis
FPC: first principle component
DHA: Docosahexaenoic acid
EPA: Eicosapentaenoic acid
TL: top-left
BR: bottom-right
BL: bottom-left
TR: top-right
PC: phosphatidylcholine
CNS: central nervous system
GPCR: G-protein coupled receptor
BBB: blood brain barrier

## Acknowledgement statement

Metabolomics data used in preparation of this article were generated by the Alzheimer’s Disease Metabolomics Consortium (ADMC). Metabolomics data will be available via the AD Knowledge Portal hosted by Sage Bionetworks. As such, the investigators within the ADMC other than named authors provided data but did not participate in analysis or writing of this report. A complete listing of ADMC investigators can be found at: https://sites.duke.edu/adnimetab/team/. The NIA supported the ADMC which is a part of NIA’s national initiatives AMP-AD and M2OVE-AD (R01AG046171, 1RF1AG059093, 1RF1AG058942, 1RF1AG051550, 3U01AG061359, 3U01AG024904-09S4, 1R01AG081322). Additional support came from 5U01AG046139, INS-273172-03, ISB-281091-03, R01AG062514, R56AG084128, RF1AG055549,

