## Supplemental Figures for "Multiomic Evidence for a Unified Model of Alzheimer’s Disease Etiology Linking Microglial Flux Capacity and Astrocyte-Neuron Metabolic Breakdown"

### Supplemental Table 1

#### Supplemental Table 1A

| Cluster 1 |
|---|
| phospholipids 18.2\|16.0 |
| CER.D18.1.16.0. |
| CER.D18.1.26.0. |
| CER.D18.2.14.0. |
| CER.D18.2.16.0. |
| CER.D18.2.26.0. |
| HEXCER.D16.1.22.0. |
| HEXCER.D16.1.24.0. |
| HEXCER.D18.2.18.0. |
| HEXCER.D18.2.20.0. |
| HEX2CER.D16.1.16.0. |
| SM.D16.1.19.0. |
| SM.D16.1.23.0..SM.D17.1.22.0. |
| SM.D16.1.24.1. |
| SM.D17.1.16.0. |
| SM.D18.0.14.0. |
| SM.D18.1.14.0..SM.D16.1.16.0. |
| SM.D18.1.16.0. |
| SM.D18.1.18.0..SM.D16.1.20.0. |
| SM.D18.1.20.0..SM.D16.1.22.0. |
| SM.D18.1.22.0..SM.D16.1.24.0. |
| SM.D18.1.23.0..SM.D17.1.24.0. |
| SM.D18.1.24.0. |
| SM.D18.1.24.1. |
| SM.D18.2.14.0. |
| SM.D18.2.16.0. |
| SM.D18.2.17.0. |
| SM.D18.2.18.0. |
| SM.D18.2.18.1. |
| SM.D18.2.20.0. |
| SM.D18.2.22.0. |
| SM.D18.2.23.0. |
| SM.D18.2.24.0. |
| PC.15.MHDA_18.2. |
| PC.16.0_16.0. |
| PC.16.0_18.0. |
| PC.16.1_18.2. |
| PC.17.0_18.2. |
| PC.17.1_18.2. |
| PC.O.16.0.16.0. |
| PC.O.18.0.18.1. |
| PC.P.16.0.14.0. |
| PC.P.16.0.16.1. |
| PC.P.16.0.18.0. |
| PC.P.16.0.18.1. |
| PC.P.18.0.22.5. |
| CE.16.0. |
| CE.18.1. |

| Cluster 2 |
|---|
| hexceramides\|phospholipids 18.1 |
| HEXCER.D18.1.16.0. |
| HEXCER.D18.1.18.0. |
| HEXCER.D18.1.20.0. |
| HEXCER.D18.1.22.0. |
| HEXCER.D18.1.24.0. |
| HEXCER.D18.1.24.1. |
| HEXCER.D18.2.22.0. |
| HEXCER.D18.2.24.0. |
| HEX2CER.D16.1.24.1. |
| HEX2CER.D18.1.16.0. |
| HEX2CER.D18.1.20.0. |
| HEX2CER.D18.1.22.0. |
| HEX2CER.D18.1.24.0. |
| HEX2CER.D18.1.24.1. |
| HEX2CER.D18.2.16.0. |
| HEX2CER.D18.2.24.1. |
| HEX3CER.D18.1.16.0. |
| HEX3CER.D18.1.18.0. |
| HEX3CER.D18.1.20.0. |
| HEX3CER.D18.1.22.0. |
| HEX3CER.D18.1.24.0. |
| HEX3CER.D18.1.24.1. |
| SM.D18.0.16.0. |
| PC.18.1_18.2. |
| PC.18.2_18.2. |
| PC.20.0_20.4. |
| PC.O.18.0.18.2. |
| PC.O.18.1.18.1. |
| PC.O.18.1.18.2. |
| PC.P.16.0.16.0. |
| PC.P.16.0.18.2. |
| PC.P.16.0.18.3. |
| PC.P.18.0.18.2. |
| PC.P.18.1.18.1. |
| PE.O.18.1.18.2. |
| PE.P.18.1.18.2...A. |
| PE.P.18.1.18.3. |
| PE.P.20.0.18.2. |
| PI.18.1_18.2. |
| CE.18.2. |
| CE.20.4. |

| Cluster 3 |
|---|
| phospholipids\|plasmalogens 20.4\|16.0 |
| PC.O.16.0.20.3. |
| PC.O.16.0.20.4. |
| PC.O.18.0.20.4. |
| PC.P.15.0.20.4...A. |
| PC.P.15.0.20.4...B. |
| PC.P.16.0.20.4. |
| PC.P.17.0.20.4...A. |
| PC.P.17.0.20.4...B. |
| PC.P.18.0.20.4. |
| PC.P.20.0.20.4. |
| PE.O.16.0.18.2. |
| PE.O.16.0.20.3. |
| PE.O.16.0.20.4. |
| PE.O.16.0.22.4. |
| PE.O.18.0.20.4. |
| PE.O.18.0.22.5. |
| PE.P.15.0.20.4...A. |
| PE.P.15.0.20.4...B. |
| PE.P.16.0.18.1. |
| PE.P.16.0.18.2. |
| PE.P.16.0.18.3. |
| PE.P.16.0.20.3...A. |
| PE.P.16.0.20.3...B. |
| PE.P.16.0.20.4. |
| PE.P.16.0.22.4. |
| PE.P.16.0.22.5...N3. |
| PE.P.16.0.22.5...N6. |
| PE.P.17.0.20.4...A. |
| PE.P.17.0.20.4...B. |
| PE.P.18.0.18.1. |
| PE.P.18.0.18.2. |
| PE.P.18.0.18.3. |
| PE.P.18.0.20.3...A. |
| PE.P.18.0.20.3...B. |
| PE.P.18.0.20.4. |
| PE.P.18.0.22.4. |
| PE.P.18.0.22.5...N3. |
| PE.P.18.0.22.5...N6. |
| PE.P.18.1.18.1...A. |
| PE.P.18.1.18.1...B. |
| PE.P.18.1.18.2...B. |
| PE.P.18.1.20.3...A. |
| PE.P.18.1.20.3...B. |
| PE.P.18.1.20.4...A. |
| PE.P.18.1.20.4...B. |
| PE.P.18.1.22.4. |
| PE.P.18.1.22.5...A. |
| PE.P.18.1.22.5...B. |
| PE.P.19.0.20.4...A. |
| PE.P.19.0.20.4...B. |
| PE.P.20.0.18.1. |
| PE.P.20.0.20.4. |
| PE.P.20.1.20.4. |
| TG.O.50.1...NL.18.1. |
| TG.O.50.2...NL.18.2. |
| TG.O.50.3...NL.18.2. |
| TG.O.52.2...NL.18.1. |
| TG.O.54.3...NL.18.1. |
| TG.O.54.4...NL.18.2. |

| Cluster 4 |
|---|
| ceramides 16.1\|18.1 |
| S1P.D16.1. |
| CER.D16.1.16.0. |
| CER.D16.1.18.0. |
| CER.D16.1.20.0. |
| CER.D16.1.22.0. |
| CER.D16.1.23.0. |
| CER.D16.1.24.0. |
| CER.D16.1.24.1. |
| CER.D17.1.16.0. |
| CER.D17.1.18.0. |
| CER.D18.1.14.0. |
| CER.D18.1.18.0. |
| CER.D18.1.19.0. |
| CER.D18.1.20.0. |
| CER.D18.1.21.0. |
| CER.D18.1.22.0. |
| CER.D18.1.23.0. |
| CER.D18.1.24.1. |
| CER.D18.2.18.0. |
| CER.D18.2.20.0. |
| CER.D18.2.21.0. |
| CER.D18.2.22.0. |
| CER.D18.2.23.0. |
| CER.D18.2.24.1. |
| CER.D19.1.16.0. |
| CER.D19.1.18.0. |
| HEXCER.D16.1.18.0. |
| HEXCER.D16.1.20.0. |
| PC.15.MHDA_18.1. |
| PC.15.MHDA_20.4. |
| PC.15.0_20.4. |
| PC.17.0_20.4. |
| PE.15.MHDA_22.6. |
| CE.20.4....OH. |

| Cluster 5 |
|---|
| acylcarnitines\|triglycerides 20.4\|18.0 |
| S1P.D18.0. |
| S1P.D18.1. |
| S1P.D18.2. |
| SM.D18.0.22.0. |
| PC.16.0_20.4. |
| PC.18.0_20.4. |
| PI.18.0_20.3...A. |
| PI.20.0_20.4. |
| AC.16.0. |
| AC.16.1. |
| AC.18.0. |
| AC.18.1. |
| AC.18.2. |
| AC.20.4. |
| AC.16.0..OH |
| AC.16.1..OH |
| AC.18.0..OH |
| AC.18.1..OH |
| TG.52.3...NL.16.1. |
| TG.52.3...NL.18.2. |
| TG.52.4...NL.16.1. |
| TG.52.4...NL.18.2. |
| TG.54.3...NL.18.1. |
| TG.54.3...NL.18.2. |
| TG.54.4...NL.18.2. |
| TG.56.6...NL.20.4. |
| TG.56.7...NL.20.4. |
| TG.56.8...NL.20.4. |

| Cluster 6 |
|---|
| phospholipids 18.0\|16.0 |
| PC.14.0_20.4. |
| PC.16.0_18.3...B. |
| PC.16.0_20.3...B. |
| PC.16.1_20.4. |
| PC.18.0_20.3. |
| PC.18.0_22.4. |
| PC.18.0_22.5...N6. |
| PE.15.MHDA_18.1. |
| PE.15.MHDA_18.2. |
| PE.15.MHDA_20.4. |
| PE.16.0_16.0. |
| PE.16.0_16.1. |
| PE.16.0_18.1. |
| PE.16.0_18.2. |
| PE.16.0_18.3...A. |
| PE.16.0_18.3...B. |
| PE.16.0_20.3. |
| PE.16.0_20.4. |
| PE.16.1_18.2. |
| PE.16.1_20.4. |
| PE.17.0_18.1. |
| PE.17.0_18.2. |
| PE.17.0_20.4. |
| PE.18.0_18.1. |
| PE.18.0_18.2. |
| PE.18.0_20.3...A. |
| PE.18.0_20.3...B. |
| PE.18.0_20.4. |
| PE.18.0_22.4. |
| PE.18.0_22.5...N3. |
| PE.18.0_22.5...N6. |
| PE.18.1_18.1. |
| PE.18.1_18.2. |
| PE.20.0_20.4. |
| PI.15.MHDA_20.4..PI.17.0_20.4. |
| PI.16.0_16.1. |
| PI.16.0_20.3...B. |
| PI.16.0_20.4. |
| PI.18.0_20.3...B. |
| PI.18.0_20.4. |
| PI.18.0_22.4. |

| Cluster 7 |
|---|
| phospholipids\|plasmalogens 22.6 |
| SM.D18.1.17.0..SM.D17.1.18.0. |
| PC.15.MHDA_22.6. |
| PC.15.0_22.6. |
| PC.16.0_22.6. |
| PC.16.1_22.6. |
| PC.17.0_22.6. |
| PC.18.1_22.6...A. |
| PC.18.1_22.6...B. |
| PC.18.2_20.5. |
| PC.O.16.0.22.6. |
| PC.O.18.0.22.6. |
| PC.P.16.0.20.5. |
| PC.P.16.0.22.6. |
| PC.P.18.0.22.6. |
| PC.P.18.1.22.6. |
| PE.O.16.0.22.6. |
| PE.O.18.0.22.6. |
| PE.O.18.1.22.6. |
| PE.P.15.0.22.6...A. |
| PE.P.15.0.22.6...B. |
| PE.P.16.0.20.5. |
| PE.P.16.0.22.6. |
| PE.P.17.0.22.6...A. |
| PE.P.17.0.22.6...B. |
| PE.P.18.0.20.5. |
| PE.P.18.0.22.6. |
| PE.P.18.1.20.5...A. |
| PE.P.18.1.20.5...B. |
| PE.P.18.1.22.6...A. |
| PE.P.18.1.22.6...B. |
| PE.P.20.0.22.6. |
| PE.P.20.1.22.6. |
| CE.20.5. |
| CE.22.6. |
| CE.22.6....OH. |

| Cluster 8 |
|---|
| triglycerides\|phospholipids 22.6 |
| PC.14.0_22.6. |
| PC.16.0_20.5. |
| PC.18.0_22.5...N3..PC.20.1_20.4. |
| PC.18.0_22.6. |
| PE.16.0_20.5. |
| PE.16.0_22.6. |
| PE.17.0_22.6. |
| PE.18.0_22.6. |
| PE.18.1_22.6...A. |
| PE.18.1_22.6...B. |
| PI.18.0_22.5...N3. |
| PI.18.0_22.6. |
| AC.20.5. |
| AC.22.6. |
| TG.52.5...NL.20.5. |
| TG.54.6...NL.20.5. |
| TG.54.6...NL.22.6. |
| TG.54.7...NL.20.5. |
| TG.54.7...NL.22.6. |
| TG.56.7...NL.20.5. |
| TG.56.7...NL.22.6. |
| TG.56.8...NL.20.5. |
| TG.56.8...NL.22.6. |
| TG.56.9...NL.22.6. |
| TG.58.10...NL.22.6. |
| TG.58.8...NL.22.6. |
| TG.58.9...NL.22.6. |

| Cluster 9 |
|---|
| phospholipids 16.0\|18.1 |
| CER.D18.1.24.0. |
| CER.D18.2.24.0. |
| PC.14.0_16.0. |
| PC.16.0_18.1. |
| PC.16.0_18.2. |
| PC.16.0_18.3...A. |
| PC.16.0_20.3...A. |
| PC.17.0_18.1. |
| PC.18.0_18.1. |
| PC.18.0_18.2. |
| PC.18.1_18.1. |
| PC.18.1_20.3. |
| PI.15.MHDA_18.1..PI.17.0_18.1. |
| PI.15.MHDA_18.2..PI.17.0_18.2. |
| PI.16.0_20.3...A. |
| PI.16.0.16.0. |
| PI.18.0_18.1. |
| PI.18.0_20.2. |
| PI.18.0_22.5...N6. |
| CE.16.1. |
| CE.18.0. |
| TG.O.50.1...NL.16.0. |
| TG.O.50.2...NL.18.1. |
| TG.O.52.0...NL.16.0. |
| TG.O.52.1...NL.16.0. |
| TG.O.52.1...NL.18.1. |
| TG.O.52.2...NL.16.0. |
| TG.O.54.2...NL.18.1. |
| CE.18.2....OH. |

| Cluster 10 |
|---|
| triglycerides 18.1\|18.0 |
| CER.M18.0.20.0. |
| CER.M18.0.22.0. |
| CER.M18.0.23.0. |
| CER.M18.0.24.0. |
| CER.M18.0.24.1. |
| CER.M18.1.18.0. |
| CER.M18.1.20.0. |
| CER.M18.1.22.0. |
| CER.M18.1.23.0. |
| CER.M18.1.24.0. |
| CER.M18.1.24.1. |
| TG.48.0...NL.16.0. |
| TG.48.0...NL.18.0. |
| TG.48.1...NL.16.1. |
| TG.48.1...NL.18.1. |
| TG.48.2...NL.16.1. |
| TG.48.2...NL.18.2. |
| TG.48.3...NL.16.1. |
| TG.49.1...NL.16.1. |
| TG.50.0...NL.18.0. |
| TG.50.1...NL.16.0. |
| TG.50.1...NL.18.1. |
| TG.50.2...NL.16.1. |
| TG.50.2...NL.18.1. |
| TG.50.2...NL.18.2. |
| TG.50.3...NL.16.1. |
| TG.50.3...NL.18.2. |
| TG.50.4...NL.20.4. |
| TG.51.0...NL.16.0. |
| TG.52.1...NL.18.0. |
| TG.52.1...NL.18.1. |
| TG.52.2...NL.16.0. |
| TG.52.2...NL.18.2. |
| TG.52.5...NL.20.4. |
| TG.53.2...NL.18.1. |
| TG.54.0...NL.18.0. |
| TG.54.1...NL.18.1. |
| TG.54.2...NL.18.0. |
| TG.54.5...NL.20.4. |
| TG.54.6...NL.20.4. |

#### Supplemental Table 1B

| Cluster 1 | Cluster 2 | Cluster 3 |
|---|---|---|
| YWHAB | ALB | HBA1 |
| YWHAZ | APOA4 | HBB |
| GAA | APOC1 | |
| ALDOA | APOC2 | |
| APOE | CP | |
| CALM2 | C9 | |
| CD44 | KNG1 | |
| CHI3L1 | PGLYRP2 | |
| DDAH1 | PON1 | |
| DKK3 | THRB | |
| ENO1 | | |
| GAPDH | | |
| GSN | | |
| GMFB | | |
| GDA | | |
| PKM | | |
| LDHB | | |
| LDHC | | |
| MDH1 | | |
| NPPA | | |
| NCAM1 | | |
| NPTX2 | | |
| NPTXR | | |
| NRXN1 | | |
| OMGP | | |
| SPP1 | | |
| PARK7 | | |
| PEBP1 | | |
| PKM | | |
| PPIA | | |
| PTPRZ1 | | |
| SCG2 | | |
| SMOC1 | | |
| SOD1 | | |
| TPI1 | | |
| VGF | | |

#### Supplemental Table 1C

| Complement/HDL | Hemoglobin | Astrogliosis | ANLS/Glycolysis | Hypoxia | Synapse | Phagocytosis |
|---|---|---|---|---|---|---|
| ALB | HBA1 | YWHAB | LDHB | PARK7 | NRXN1 | GSN |
| APOA4 | HBB | YWHAZ | LDHC | PPIA | NCAM1 | SPP1 |
| APOC1 | | CHI3L1 | ALDOA | | NPTX2 | PEBP1 |
| APOC2 | | CD44 | PKM | | NPTXR | |
| CP | | | PKM | | | |
| C9 | | | MDH1 | | | |
| KNG1 | | | GAA | | | |
| PGLYRP2 | | | ENO1 | | | |
| PON1 | | | | | | |
| THRB | | | | | | |

**Supplemental Table 1:** Results of plasma lipidomic **(1A)** and CSF mass spec protein **(1B)** clustering based on Wolfram Mathematica clustering algorithm over 602 ADNI subjects for whom both data types were available (all 4 quartiles). **1C)**: A subset of protein cluster 1 in panel B was manually separated into 5 functional groups.

### Supplemental Figure 1

#### Supplemental Figure 1A

| Quadrant | Subjects | Mean APOE4 allele copies | Fraction Female | Median Age |
|---|---|---|---|---|
| TL | 63 | 0.444444 | 0.888889 | 68.2 |
| TR | 13 | 0.0769231 | 0.538462 | 67.9 |
| BR | 37 | 0.405405 | 0.459459 | 69.0 |
| BL | 51 | 0.372549 | 0.705882 | 69.3 |
| L | 114 | 0.412281 | 0.807018 | 68.95 |
| R | 50 | 0.32 | 0.48 | 68.95 |
| T | 76 | 0.381579 | 0.828947 | 68.1 |
| B | 88 | 0.386364 | 0.602273 | 69.3 |

#### Supplemental Figure 1B

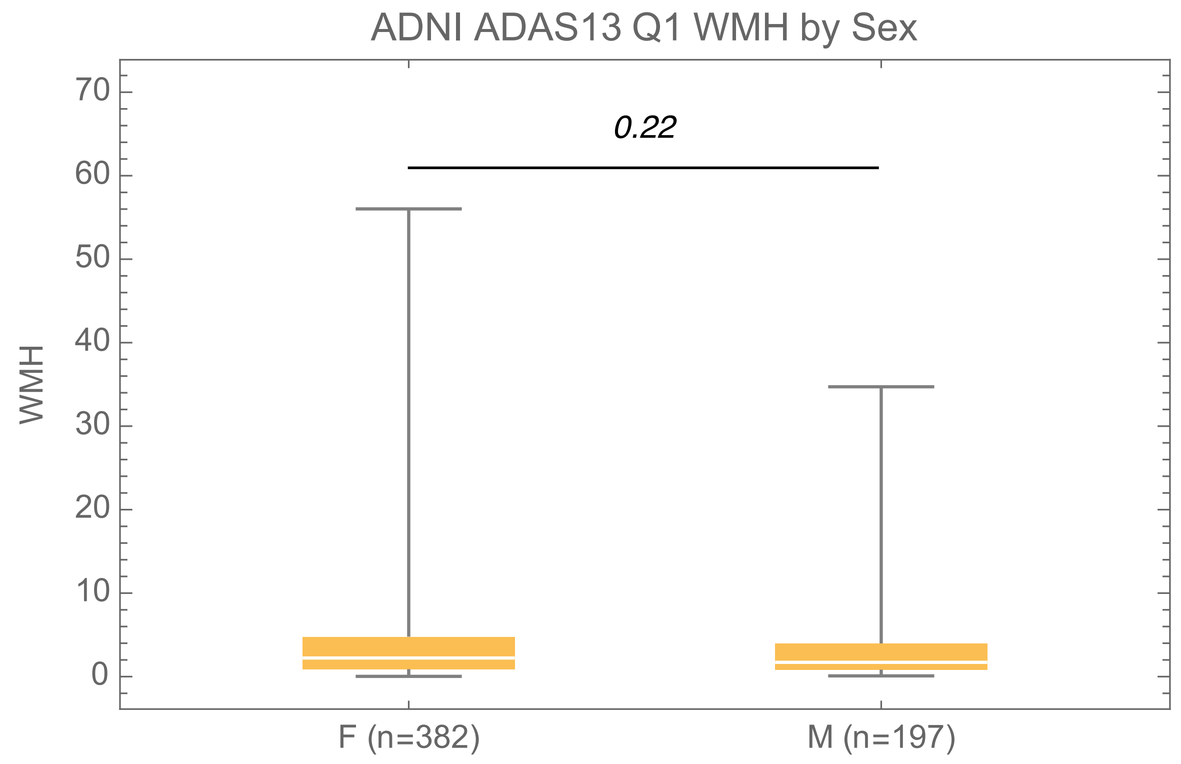


**Supplemental Figure 1A:** Q1 distribution of sex and WMH ( TL = 89%, 56/63).
**1B:** WMH for all ADNI individuals who fell within the ADAS13 range for Q1

### Supplemental Table 2

#### Supplemental Table 2A

| | | | | |
|---|---|---|---|---|
| APOM | AHSG | IGHA1\|IGHA2 | BPIFB1 | DDR1 |
| GDA | SERPINA10 | CFD | ERP29 | FCN3 |
| C8G | KNG1 | SAA4 | GLIPR2 | INHBC |
| CFHR5 | MYOC | A1BG | F13A1\|F13B | ALB |
| SGTB | PSMB9 | HABP2 | RCAN3 | PLA2G4A |
| AOC3 | CDKL2 | RHOQ | F11 | RAD51C |
| MAPK6 | DKK4 | SIRT2 | C5 | APCS |
| C3 | SHH | APOA1 | LCN2 | GHR |
| CFP | SERPINF2 | CCL18 | LBP | F10 |
| ADAMTS13 | FCGR1A | CPB2 | SERPINA1 | PLG |
| C6 | CFB | KLKB1 | IL1RL1 | PKM |
| MST1 | C5\|C6 | ITIH4 | F9 | CCL16 |
| HRG | MAP2K4 | F13B | SCGB3A1 | CPN2 |
| C14orf93 | ASPN | FBP1 | ITIH1 | AZGP1 |
| C1RL | C4BPA | | | |

#### Supplemental Table 2B

| Term | Overlap | P-value | Adjusted P-value | Odds Ratio | Combined Score | Genes |
|---|---|---|---|---|---|---|
| **Complement and coagulation cascades** | 18/70 | 2.07E-29 | 6.2E-27 | 127.4 | 8415.4 | CFD;CPB2;SERPINA1;F10;SERPINF2;F11;F13A1;C4BPA;PLG;KNG1;C3;C5;C8G;F9;C6;F13B;KLKB1;CFB |
| **Alternative complement pathway** | 7/11 | 1.89E-15 | 2.8E-13 | 536.4 | 18184.7 | C3;CFD;C5;C6;C8G;CFP;CFB |
| **Hemostasis pathway** | 16/468 | 7.74E-12 | 7.8E-10 | 12.3 | 315.0 | CFD;SERPINA1;F10;SERPINF2;F11;F13A1;PLA2G4A;APOA1;PLG;KNG1;RAD51C;F9;ALB;F13B;HRG;KLKB1 |
| **Response to elevated platelet cytosolic calcium** | 9/83 | 1.61E-11 | 1.1E-09 | 38.3 | 952.6 | CFD;SERPINA1;SERPINF2;ALB;F13A1;APOA1;PLG;HRG;KNG1 |
| **Fibrin clot formation (clotting cascade)** | 7/32 | 1.82E-11 | 1.1E-09 | 85.7 | 2120.2 | F9;F10;F11;F13A1;F13B;KLKB1;KNG1 |
| **Blood clotting cascade** | 6/23 | 1.69E-10 | 8.5E-09 | 106.5 | 2395.7 | F9;F10;F11;SERPINF2;PLG;F13B |
| **Coagulation intrinsic pathway** | 5/15 | 1.53E-09 | 6.6E-08 | 148.6 | 3016.9 | F9;F10;F11;KLKB1;KNG1 |
| **Platelet activation, signaling and aggregation** | 10/205 | 3.17E-09 | 1.2E-07 | 16.3 | 319.4 | CFD;SERPINA1;SERPINF2;ALB;F13A1;PLA2G4A;APOA1;PLG;HRG;KNG1 |
| **Complement cascade** | 7/77 | 1.14E-08 | 3.8E-07 | 30.6 | 558.7 | CFD;C3;C5;C8G;C6;C4BPA;CFB |
| **Prothrombin activation intrinsic pathway** | 5/32 | 9.80E-08 | 3.0E-06 | 55.0 | 887.7 | F9;F10;F11;KLKB1;KNG1 |

**Supplemental Table 2A:** Somascan proteins from Q1 (L vs R quadrants) used for EnrichR analysis **(2B)**.

### Supplemental Table 3

#### Supplemental Table 3A

| | | | | |
|---|---|---|---|---|
| NEFL | AHSG | TYMP | VTN | C7 |
| FCN3 | INHBC | PPBP | HMOX1 | SNX3 |
| HABP2 | PDE4C | RCAN3 | EIF1 | NECAP2 |
| CHMP4A | ADGRD1 | UBE2D3\|UBB | PDLIM3 | RAD51C |
| DKK4 | SIRT2 | TPD52L2 | SNX15 | CCN3 |
| IL16 | AGRP | CFP | PRSS1 | LBP |
| F7 | CD5L | REN | SELE | SLPI |
| SFN | CCL16 | DBNL | CD27 | PAM |
| LILRA3 | MGP | PLOD2 | ACP7 | ARSK |
| PENK | NMB | G3BP2 | NEFH | |

#### Supplemental Table 3B

| Term | Overlap | P-value | Adjusted P-value | Old Adjusted P-value | Odds Ratio | Combined Score | Genes |
|---|---|---|---|---|---|---|---|
| Alternative complement pathway | 2/11 | 3.32E-04 | 8.2E-02 | 0.0E+00 | 92.3 | 739.5 | C7;CFP |
| Peptide G-protein coupled receptors | 4/192 | 1.34E-03 | 8.9E-02 | 0.0E+00 | 9.1 | 60.4 | NMB;PENK;PPBP;CCL16 |
| Downregulation of SMAD2/3-SMAD4 transcriptional activity | 2/23 | 1.50E-03 | 8.9E-02 | 0.0E+00 | 39.5 | 257.2 | UBB;UBE2D3 |
| Cellular response to hypoxia | 2/25 | 1.77E-03 | 8.9E-02 | 0.0E+00 | 36.1 | 228.7 | UBB;UBE2D3 |
| PTM: gamma carboxylation, hypusine formation and arylsulfatase activation | 2/28 | 2.22E-03 | 8.9E-02 | 0.0E+00 | 31.9 | 195.1 | F7;ARSK |
| Endosomal sorting complex required for transport (ESCRT) pathway | 2/28 | 2.22E-03 | 8.9E-02 | 0.0E+00 | 31.9 | 195.1 | UBB;CHMP4A |
| Leptin influence on immune response | 3/110 | 2.63E-03 | 8.9E-02 | 0.0E+00 | 11.8 | 70.3 | VTN;IL16;PPBP |
| Negative regulators of RIG-I/MDA5 signaling | 2/33 | 3.08E-03 | 8.9E-02 | 0.0E+00 | 26.8 | 154.8 | UBB;UBE2D3 |
| Interleukin-7 interactions in immune response | 2/34 | 3.27E-03 | 8.9E-02 | 0.0E+00 | 25.9 | 148.5 | PPBP;CCL16 |
| FRA pathway | 2/37 | 3.86E-03 | 9.5E-02 | 0.0E+00 | 23.7 | 131.8 | MGP;HMOX1 |

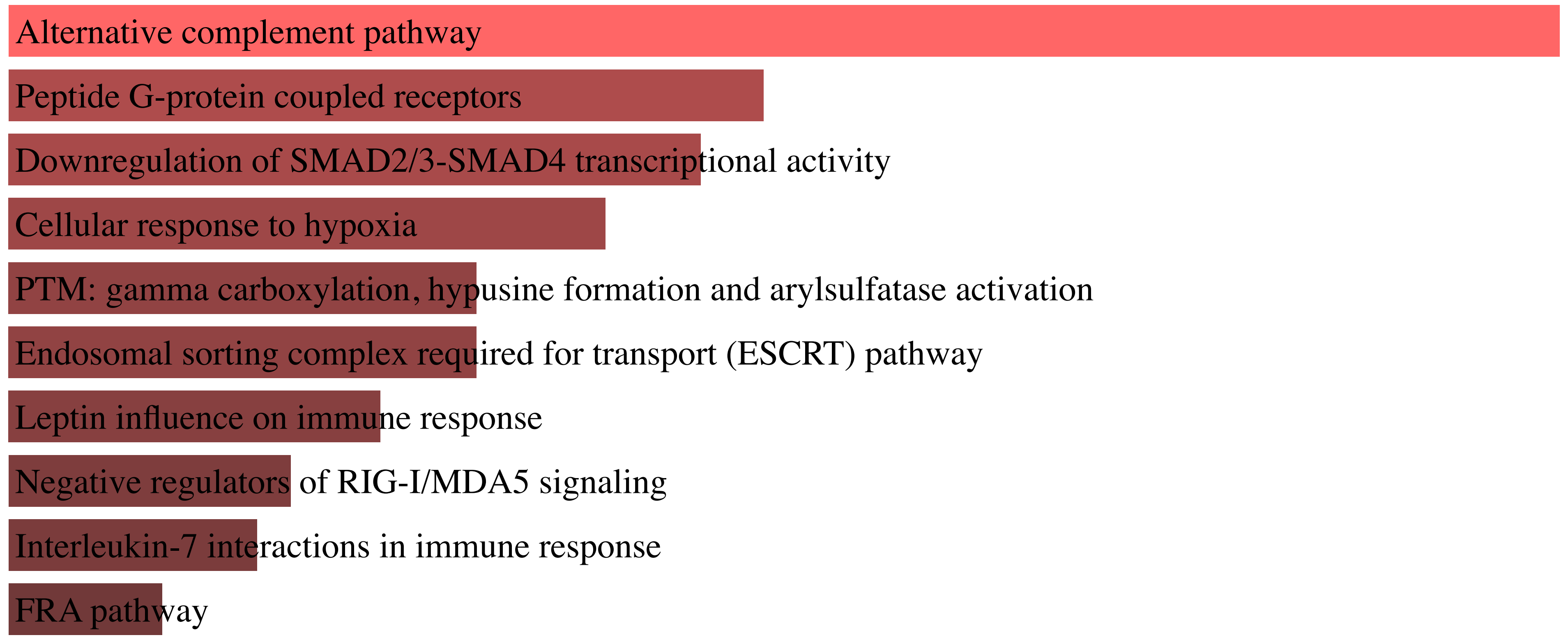


**Supplemental Table 3:** Somascan proteins from Q1 (T vs B quadrants) used for EnrichR analysis **(3B)**.

### Supplemental Figure 3

#### Supplemental Figure 3

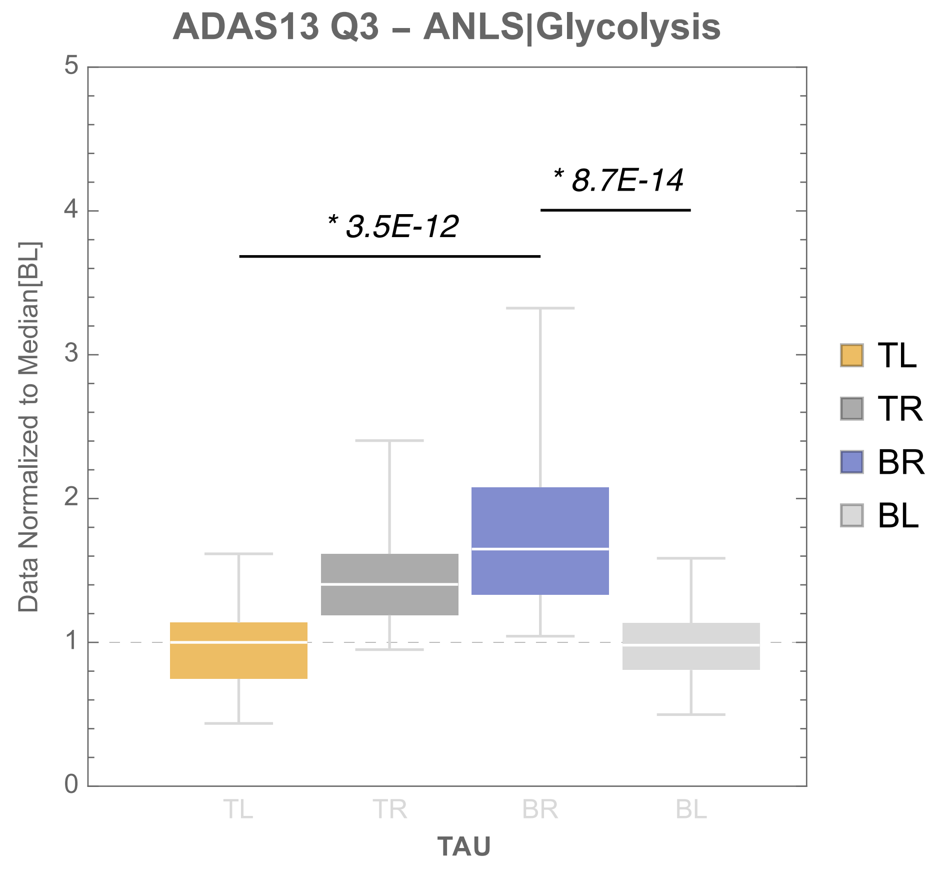


**Supplemental Figure 3:** Quantification of Tau per each quadrant in Q3

### Supplemental Table 4

#### Supplemental Table 4A

| | | | | |
|---|---|---|---|---|
| PPIH | ARL3 | EIF1B | KIFBP | WFDC10A |
| RAB21 | AKR1D1 | TCEAL8 | CYB5R2 | NDUFA5 |
| RAB5B | ALDH1A2 | IMPACT | NREP | DSG4 |
| XAGE2 | RAB7A | CRYL1 | GYG2 | INPP5A |
| TOM1 | RAC1 | FGF1 | SLC27A2 | |

#### Supplemental Table 4B

| Term | Overlap | P-value | Adjusted P-value | Odds Ratio | Combined Score | Genes |
|---|---|---|---|---|---|---|
| Bile acid and bile salt biosynthesis via 24-hydroxycholesterol | 2/10 | 6.17E-05 | 6.42E-03 | 226.9 | 2199.3 | AKR1D1;SLC27A2 |
| Bile acids and bile salt biosynthesis | 2/10 | 6.17E-05 | 6.42E-03 | 226.9 | 2199.3 | AKR1D1;SLC27A2 |
| Bile acid and bile salt biosynthesis via 7-alpha-hydroxycholesterol | 2/15 | 1.44E-04 | 9.95E-03 | 139.6 | 1235.3 | AKR1D1;SLC27A2 |
| Bile acid and bile salt metabolism | 2/27 | 4.76E-04 | 2.47E-02 | 72.5 | 555.1 | AKR1D1;SLC27A2 |
| Phagosome | 3/154 | 8.05E-04 | 3.35E-02 | 18.8 | 133.6 | RAB5B;RAC1;RAB7A |
| Post-translational regulation of adherens junction stability and disassembly | 2/48 | 1.51E-03 | 5.22E-02 | 39.4 | 256.0 | RAC1;RAB7A |
| MAPK cascade role in angiogenesis | 2/64 | 2.66E-03 | 7.90E-02 | 29.2 | 173.2 | RAC1;FGF1 |
| FGFR1b ligand binding and activation | 1/5 | 5.99E-03 | 1.13E-01 | 217.1 | 1111.1 | FGF1 |
| Amoebiasis | 2/105 | 6.99E-03 | 1.13E-01 | 17.5 | 87.1 | RAB5B;RAB7A |
| Bile acid and bile salt biosynthesis via 27-hydroxycholesterol | 1/6 | 7.18E-03 | 1.13E-01 | 173.7 | 857.3 | AKR1D1 |

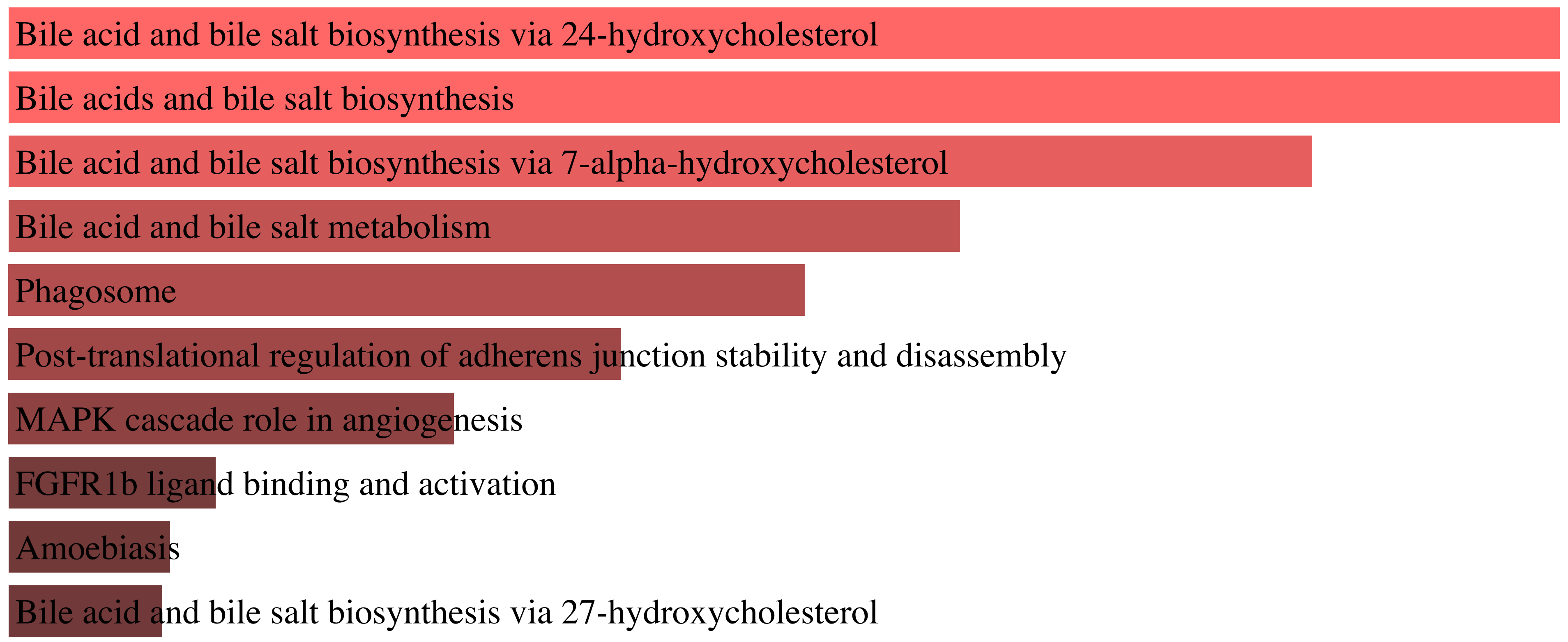


**Supplemental Table 4:** Somascan proteins from Q2 that correlate with CER.D19.1.18.0. (T vs B quadrants) used for EnrichR analysis **(4B)**.

### SNPs from AD GWAS

#### (all significance calculations based on Q2 only)

**Supplemental Figure 4A**

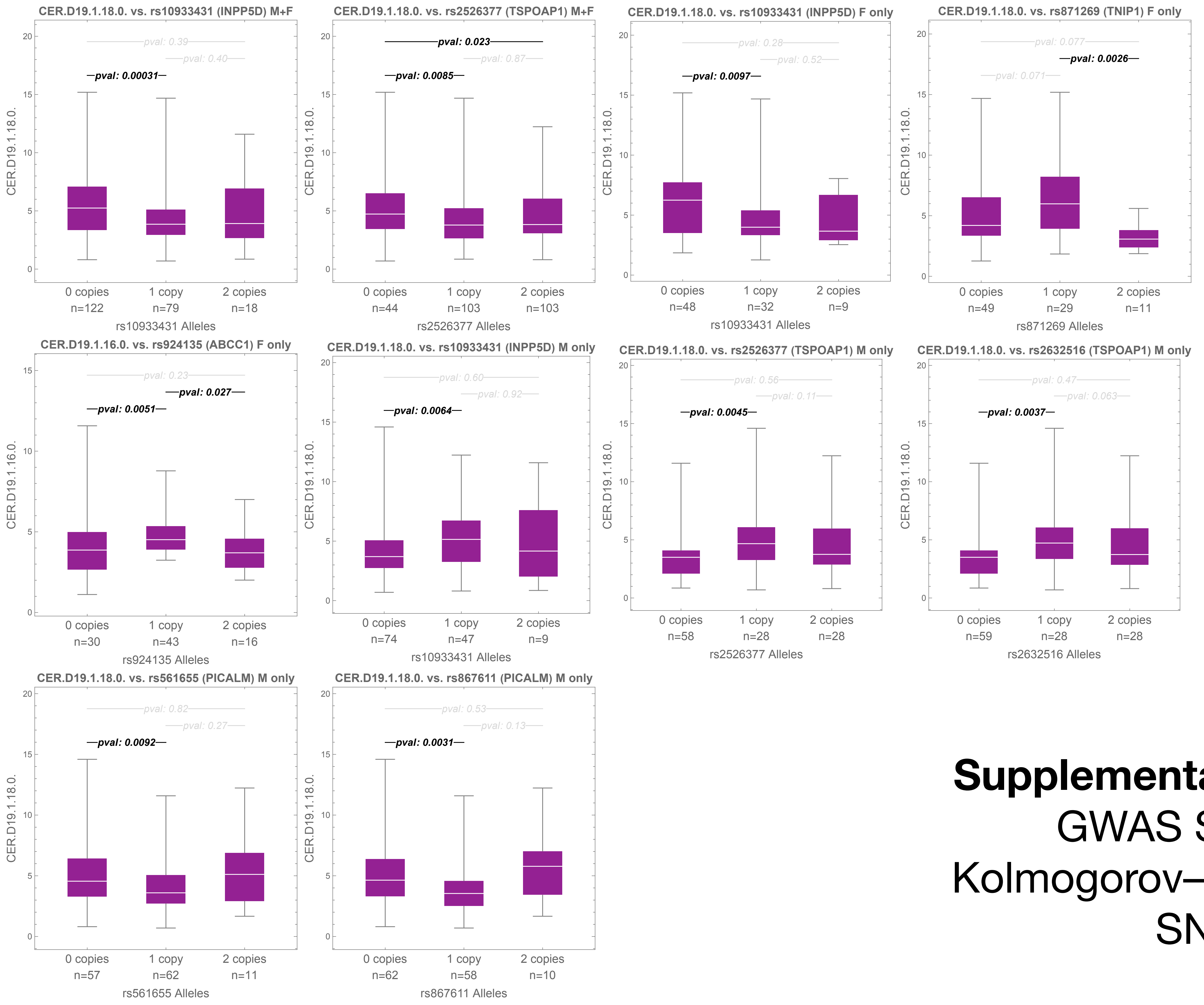


| locus | cytoband | SNP | CHR | BP | A1 | A2 | GENE | FRQ | global_maf | OR | P | study |
|---|---|---|---|---|---|---|---|---|---|---|---|---|
| 10 | 2q37.1 | rs10933431 | 2 | 233 117 202 | C | G | INPP5D | 0.2343 | 0.372197 | 0.93 | 0.00000000000000000362 | Bellenguez |
| 10 | 2q37.1 | rs10933431 | 2 | 233 981 912 | C | G | INPP5D | 0.223 | 0.372197 | 0.91 | 0.0000000034 | Kunkle |
| 10 | 2q37.1 | rs10933431 | 2 | 233 981 912 | C | G | INPPD5 | 0.244487 | 0.372197 | 0.984682 | 0.000000000762 | Jansen |
| 20 | 5q33.1 | rs871269 | 5 | 150 432 388 | C | T | TNIP1 | 0.32 | 0.393324 | 0.991951 | 0.00000000137 | Wightman |
| 20 | 5q33.1 | rs871269 | 5 | 151 052 827 | C | T | TNIP1 | 0.3264 | 0.393324 | 0.96 | 0.00000000867 | Bellenguez |
| 52 | 11q14.2 | rs867611 | 11 | 85 776 544 | A | G | PICALM | 0.317416 | 0.268237 | 0.979778 | 0.00000000000000000148 | Jansen |
| 52 | 11q14.2 | rs561655 | 11 | 85 800 279 | G | A | PICALM | 0.65 | 0.300744 | 1.01441 | 0.000000000000000000000000124 | Wightman |
| 82 | 17q22 | rs2632516 | 17 | 56 409 089 | G | C | TSPOAP1–AS1 | 0.46 | 0.495851 | 0.993822 | 0.000000000746 | Wightman |
| 82 | 17q23.2 | rs2526377 | 17 | 58 332 680 | A | G | TSPOAP1 | 0.4449 | 0.496494 | 0.95 | 0.00000000000158 | Bellenguez |

**Supplemental Figure 4A:** CER.D19.1.18.0. levels for AD GWAS SNPs found to be significant based on Kolmogorov–Smirnov statistics (168 AD summary GWAS SNPs considered), grouped by sex.

### Other SNPs

#### (all significance calculations based on Q2 only or Q23 as labeled)

**Supplemental Figure 4B** **SPTLC3, Q2** **NR1H3 (LXRA), Q23**

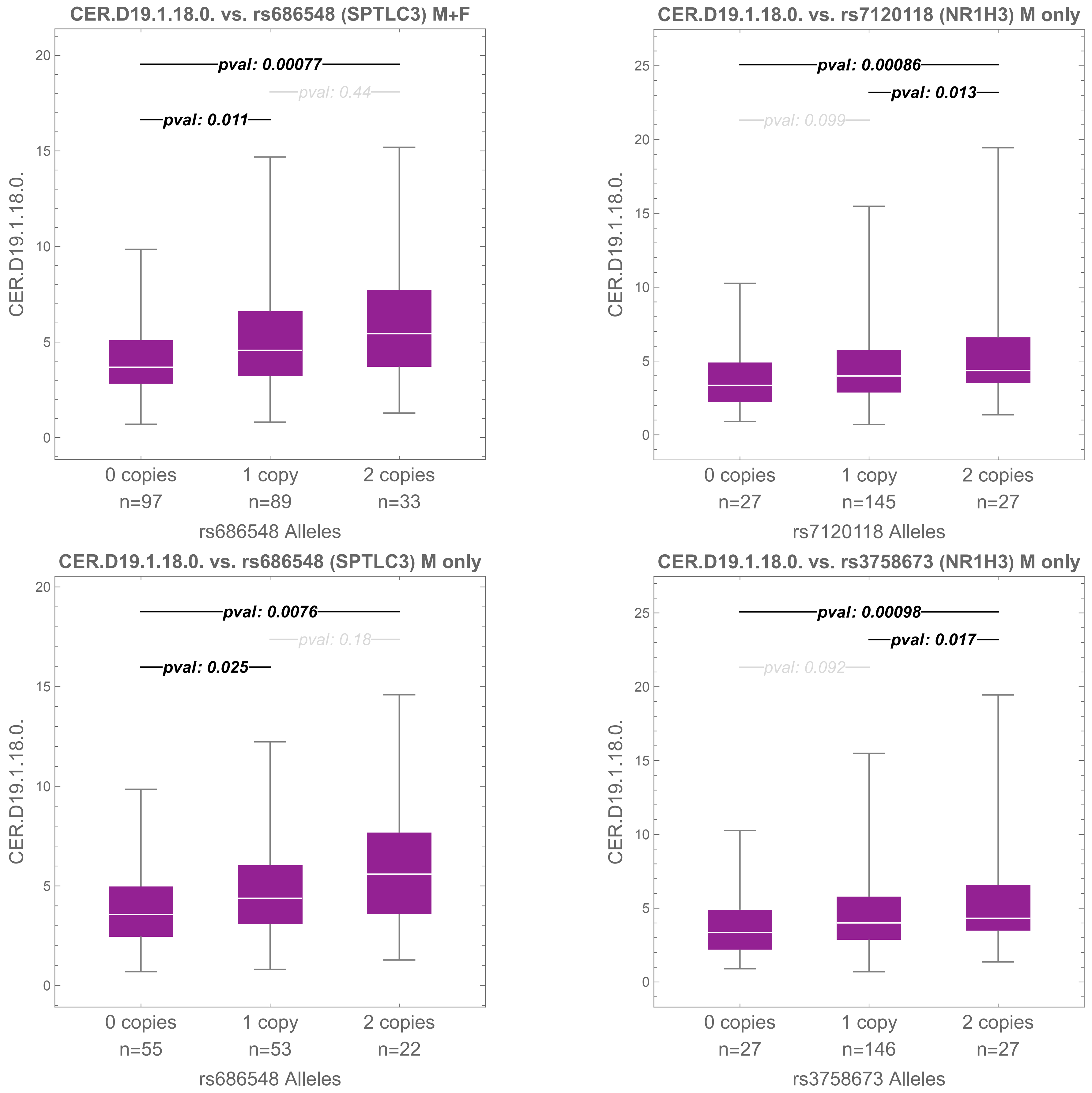


**Supplemental Figure 4B:** CER.D19.1.18.0. levels for SPTLC3 and NR1H3 (LXRA) found to be significant based on Kolmogorov–Smirnov statistics (168 AD summary GWAS SNPs considered), grouped by sex.

### Supplemental Figure 5

#### **Supplemental Figure 5**

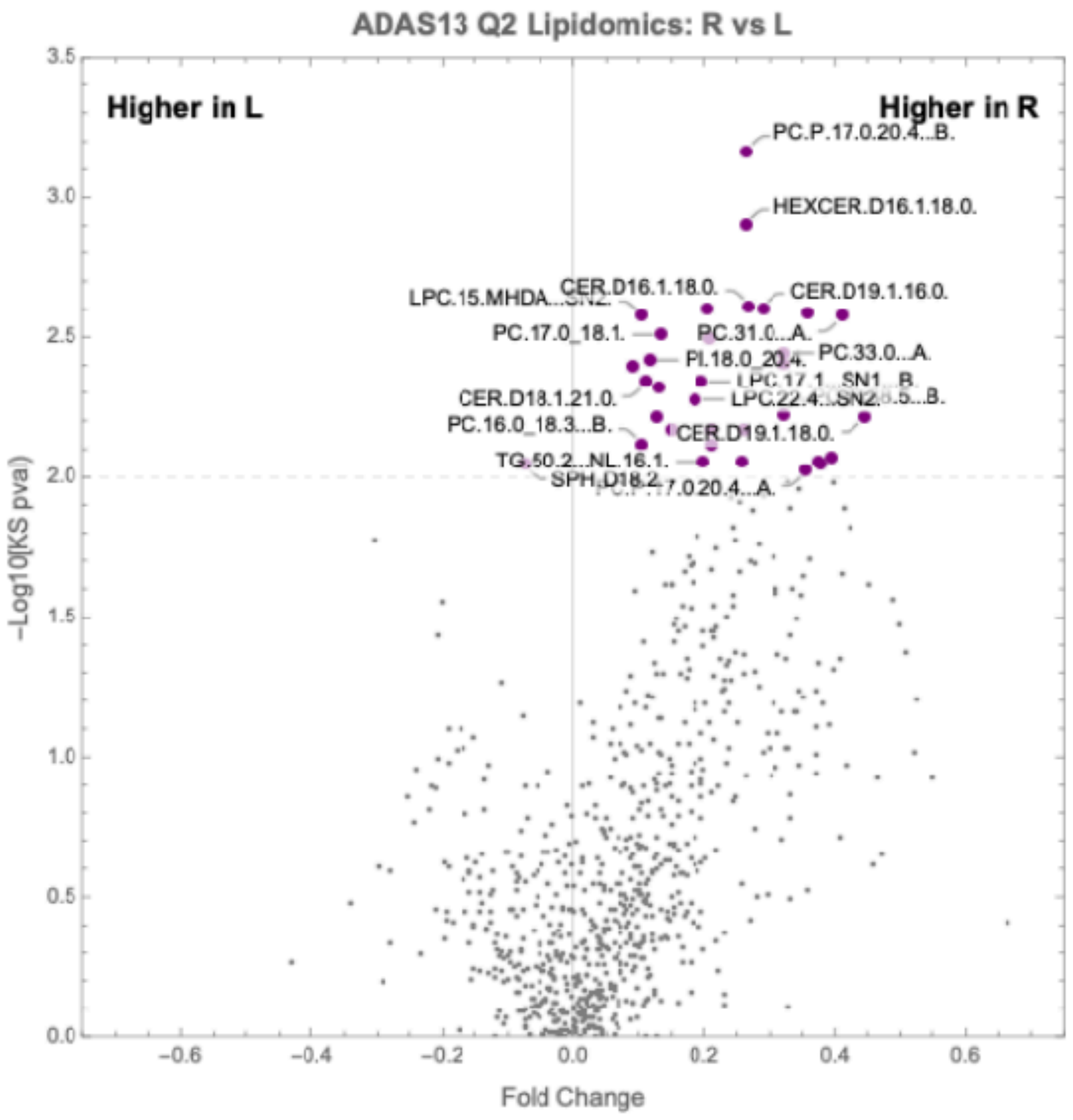


**Supplemental Figure 5:** Volcano plot for Q2 individuals (L vs R) lipidomics. P-value cutoff = 0.01. Fold change cutoff = 0.15.
